# Easymode: general pretrained networks for cellular cryo-ET enable flexible approaches to subtomogram averaging

**DOI:** 10.64898/2026.05.19.726344

**Authors:** Mart G. F. So-Last, Alister Burt, Thomas Hale, Matteo Allegretti

## Abstract

Cellular cryo-electron tomography (cryo-ET) reveals high-resolution details of macromolecules within their native cellular environment. However, *in situ* cryo-ET datasets are large and highly heterogeneous, which makes comprehensive identification and extraction of the many different elements of cellular architecture for high-resolution analysis a challenging, time-consuming and often tedious task. Here we present *easymode*, a library of pretrained general segmentation networks for cryo-ET, trained on over 4,000 tilt series spanning a large and diverse variety of sources. Easymode enables *in situ* structural determination workflows by rendering tomogram content computationally accessible, without requiring any per-dataset training. Beyond directly facilitating high-resolution subtomogram averaging of a selection of widely prevalent complexes, we show how easymode can be used to leverage cellular context in subtomogram averaging workflows, helping identify, align, or filter particle sets, and enabling automated mapping of the cellular landscape surrounding target proteins. We use easymode to determine the *in situ* structure of rare inosine monophosphate dehydrogenase (IMPDH) filaments at 4.0 A resolution, and to map and visualize the surrounding cellular environment.

## Introduction

Cellular cryo-electron tomography (cryo-ET) enables the visualization of macromolecular complexes within their native environment, bridging structural and cell biology at subnanometer resolution^1–3^. Driven by advances in cryo-focused ion beam milling^4–6^, automated data acquisition^7–9^, and data processing^10–19^, cryo-ET has matured into a powerful tool that can address fundamental questions in cell biology. Among many examples, recent *in situ* studies have revealed details of the cellular architecture of Asgard archaea^20–23^, captured the ribosomal translation cycle in bacteria and human cells^24,25^, visualized the passage of HIV-1 capsids through nuclear pore complexes in human immune cells^26–28^, and resolved the molecular organisation of the native sarcomere^29,30^. Each of these studies represents a major contribution to the understanding of the molecular biology of these systems, and was enabled by cellular cryo-ET.

In most cryo-ET studies, structure determination efforts focus on one or a small number of specific target protein complexes. These targets are almost always situated in a cellular landscape that is visually dominated by a set of large, prevalent features: ribosomes, cytoskeletal filaments, membranous compartments, nuclear pore complexes, nucleosomes, and major organelles such as mitochondria, the nucleus, nuclear envelope, and the cytoplasm itself. Together, these features form the structural landscape in which any individual complex of interest is embedded. Mapping them can therefore reveal the biological context surrounding a target and enable a more informed interpretation of its structure *in situ*.

Underpinning both structure determination and contextual mapping is feature detection, for which a range of computational approaches exists. For particle identification, approaches include template matching^31–33^, embedding-based methods^34–36^, segmentation-based approaches^37–40^, or manual picking. For the detection of broader cellular features - organelles such as mitochondria and nuclei, membranes, granules, and other contextual structures - manual segmentation or using trained machine learning (ML) models are the most common approaches^38,41–45^. However, generating segmentations for a new dataset typically requires considerable effort, either in direct annotation or in the preparation of training data for a bespoke network.

An effective alternative is to use pretrained general networks for segmentation, which can be applied directly to new data without any per-dataset training or annotation. MemBrain^43,44^, a widely adopted tool for membrane segmentation, chiefly demonstrates the utility of this approach, and a few other pretrained models have been published for a limited set of cellular features, including microtubules, membranes, and ribosomes (TARDIS^46^, DeePiCt^38^, cryoSIAM^35^), although these models are often trained using a small number of tomograms and data sources. Consequently, the range of features and organisms for which reliable pretrained networks are currently available remains very limited in scope. If a broader set of pretrained general networks for cellular cryo-ET were available, this could substantially simplify the mapping of the cellular landscape. In turn, this would enable users to survey large datasets for specific features of interest, to build models of cellular architecture by combining segmentations of many different components, and to use the resulting biological information to curate particle selections and interpret molecular structures within their native context.

Here, we present *easymode:* a large collection of pretrained general segmentation networks for cellular cryo-ET, designed to facilitate *in situ* structure determination workflows by rendering the biological content of tomograms computationally accessible. The network collection encompasses models for the detection of common features including macromolecular complexes (ribosomes, the T-complex protein Ring Complex or TRiC, and vaults), cytoskeletal elements (microtubules, intermediate filaments, and actin), nuclear architecture (the nucleus, nuclear envelope, and nuclear pore complexes), the cytoplasm, mitochondria, membranes, granules (cytoplasmic and mitochondrial) and auxiliary features (ice particles and sample boundaries).

To assess the reliability of the easymode networks, we validated the networks by using their segmentation output to generate particle coordinates for subtomogram averaging (STA) or using standard mask-based segmentation metrics. We first show how segmentation-based particle detection with easymode can be used to support *in situ* structure determination in six examples: averaging ribosomes, microtubules, actin, TRiC, nuclear pore complexes and vault complexes, with the first four solved at subnanometer resolution. Next, we present validation results for the networks that segment pleiomorphic structures such as organelles, granules, and the sample boundaries.

As easymode is intended to be a tool to support *in situ* structure determination for contextual studies, we then demonstrate several examples of how, by combining the output of multiple pretrained segmentation networks, easymode facilitates STA. In four use-cases we show how to combine the macromolecular networks with both the organellar and boundary networks to largely improve *in situ* structural determination without the need of classification. Finally, we show how we used easymode to find and solve the *in situ* structure of *inosine monophosphate dehydrogenase* (IMPDH) filaments at 4.0 A resolution, and to build detailed maps of the cellular environment surrounding this protein assembly.

The software is released as an open-source tool and is linked to an online model repository from which the required network weights are automatically downloaded at inference time, so that users require no prior experience with machine learning or neural network training. Importantly, the model library and the training data collection have both been designed to be extensible. New models can be added to the repository at a later time and are then automatically distributed for use. Easymode also includes a voluntary reporting tool which enables users to flag model failures and upload the relevant tomograms to our data server, allowing us to improve the models over time by including additional challenging cases in the training data. The software, documentation, and an illustrated model library are available via mgflast.github.io/easymode.

## Results

### A large and diverse training data collection for cellular cryo-ET

To be able to train networks that generalize across the diversity of cellular cryo-ET data, we began by compiling a large and diverse training data collection. We downloaded tilt series from EMPIAR^47^ and the Cryo-ET Data Portal^48^ and collected contributions from colleagues at the LMB and several external groups. In total, the resulting training collection (**Table 1**) comprises 4,120 tilt series drawn from 58 distinct datasets, covering 25+ different eukaryotic and prokaryotic species and a broad range of imaging parameters including variation in detector type, pixel size, acceleration voltage, defocus, specimen thickness, tilt range, electron doses, and sample types (lysates, protein and organelle isolates, whole cells, focused ion beam (FIB)-milled cells, plasma-FIB-milled high-pressure-frozen samples). The majority of datasets feature eukaryotic species and almost half (23 out of 58) comprise human samples, spanning a range of cell types including spermatozoa, T lymphocytes, macrophages, neutrophils, fibroblasts and several commonly cultured cell lines (including HeLa, Jurkat, U2OS, HEK293T, and RPE1).

**Table 1:** Composition of the training data collection. **a)** The source of each dataset is indicated via references to the original articles, when available; absence of a reference means no publication has appeared at the time of writing. EMPIAR: EM Public Image Archive^**47**^, CDP: Cryo-ET Data Portal^**48**^, EXT: private contributions from external groups (all contributors are listed in the Acknowledgements), INT: internal contributions from groups at our institute (mostly unpublished). HPF/pFIB is shorthand for high-pressure-frozen, plasma-FIB-milled; the other lamella are mostly from gallium-FIB milled plunge-frozen cells. **b)** This column lists the pixel size of the original tilt series. All tomograms were reconstructed at a common 10.0 Å/px. **c)** This mixed contribution consists of reconstructed tomograms submitted by users. At the time of writing, the majority comprised unidentified prokaryotic species in an environmental isolate.

| Dataset ID | Species and sample type<sup>a</sup> | Tilt series (n) | Pixel size (Å)<sup>b</sup> |
| --- | --- | --- | --- |
| EMPIAR-10491 | Purified apoferritin (*H. sapiens*)<sup>11</sup> | 37 | 0.79 |
| EMPIAR-10164 | Purified HIV particles<sup>49</sup> | 10 | 0.68 |
| EMPIAR-11111 | Purified 70S ribosome (*E. coli*)<sup>7</sup> | 25 | 1.07 |
| EMPIAR-12612 | *S. oleracea* chloroplasts lamella<sup>50</sup> | 23 | 3.52 |
| EMPIAR-11830 | *C. reinhardtii* lamella<sup>42</sup> | 88 | 1.96 |
| EMPIAR-11747 | *T. pseudonana* lamella<sup>51</sup> | 7 | 1.07 |
| EMPIAR-11078 | *C. reinhardtii* lamella<sup>52</sup> | 23 | 3.42 |
| EMPIAR-10499 | *M. pneumonia* whole cells<sup>11</sup> | 65 | 1.70 |
| EMPIAR-11897 | *H. sapiens* (extracellular matrix), HPF/pFIB lamella<sup>53</sup> | 39 | 2.14 |
| EMPIAR-10493 | SARS-CoV2 purified virions<sup>54</sup> | 20 | 1.53 |
| EMPIAR-12176 | *E. intestinalis* lamella<sup>55</sup> | 24 | 2.06 |
| EMPIAR-10989 | *H. sapiens* (RPE1) whole cells (periphery)<sup>38</sup> | 3 | 3.45 |
| EMPIAR-11896 | EhV-201 purified particles<sup>56</sup> | 40 | 2.08 |
| EMPIAR-11058 | *T. kivui* lamella<sup>57</sup> | 17 | 3.52 |
| EMPIAR-11845 | *D. discoideum* lamella<sup>58</sup> | 152 | 2.18 |
| EMPIAR-11561 | *H. sapiens* (HeLa) lamella<sup>59</sup> | 15 | 3.43 |
| EMPIAR-12457 | *H. sapiens* (macrophage) lamella<sup>27</sup> | 39 | 2.41 |
| EMPIAR-12460 | *M. musculus* (embryonic stem cells) lamella<sup>60</sup> | 159 | 2.68 |
| EMPIAR-10988 | *S. pombe* lamella<sup>38</sup> | 9 | 3.37 |
| EMPIAR-11198 | *E. amylovora*, RAY phage infected, lamella<sup>61</sup> | 32 | 4.27 |
| EMPIAR-10466 | *S. cerevisiae* lamella<sup>62</sup> | 177 | 3.45 |
| EMPIAR-12425 | *H. sapiens* (HeLa) CEMOVIS<sup>63</sup> | 59 | 1.69 |
| EMPIAR-13289 | *M. musculus* (hippocampus) HPF/pFIB lamella<sup>64</sup> | 58 | 3.05 |
| EMPIAR-13281 | *M. musculus* (hippocampus) HPF/pFIB lamella<sup>64</sup> | 147 | 1.98 |
| EMPIAR-11538 | *H. sapiens* (HEK293T) lamella<sup>25</sup> | 357 | 1.19 |
| EMPIAR-13145 | *H. sapiens* (HeLa) lamella<sup>41</sup> | 239 | 2.31 |
| EMPIAR-11899 | *D. discoideum* lamella<sup>65</sup> | 207 | 1.22 |
| CDP-10004 | *C. elegans* HPF/pFIB lamella<sup>66</sup> | 100 | 1.50 |
| CDP-10452 | *M. mycoides* whole cells | 15 | 1.53 |
| CDP-10440 | *E. coli* lysate with added proteins<sup>67</sup> | 19 | 1.53 |
| CDP-10434 | *H. sapiens* (HEK293) whole cell periphery<sup>68</sup> | 20 | 2.17 |
| CDP-10431 | *H. sapiens* (HEK293) whole cell periphery<sup>68</sup> | 86 | 2.17 |
| CDP-10455 | *E. coli* whole cells inoculated with PP7 VLP<sup>69</sup> | 10 | 1.50 |
| CDP-10444 | *H. sapiens* purified endo/lysosomes | 10 | 1.54 |
| EXT-001 | *S. cerevisiae*, HPF/pFIB lamella | 64 | 1.56 |
| EXT-002 | *H. sapiens* (neutrophil) lamella<sup>70</sup> | 231 | 1.56 |
| EXT-003 | *D. melanogaster* (flight muscle) lamella | 149 | 1.18 |
| EXT-004 | *E. scolopes* + *V. fischeri* HPF/pFIB lamella<sup>71</sup> | 286 | 1.89 |
| EXT-005 | *M. musculus* (cardiomyocyte) lamella | 24 | 2.32 |
| EXT-006 | *H. sapiens* (U2OS) whole cells (periphery) | 26 | 2.74 |
| EXT-007 | *H. sapiens* (U2OS) lamella | 40 | 2.15 |
| EXT-008 | *S. rosetta* lamella | 55 | 3.10 |
| INT-001 | *H. sapiens* (HeLa) lamella | 60 | 1.56 |
| INT-002 | *H. sapiens* (spermatozoa) lamella<sup>72</sup> | 56 | 1.57 |
| INT-003 | *H. sapiens* (fibroblast) lamella | 52 | 1.33 |
| INT-004 | *M. musculus* lamella | 20 | 1.69 |
| INT-005 | *P. aeruginosa* lamella | 30 | 2.13 |
| INT-006 | Purified eukaryotic microtubule organizing center | 23 | 1.68 |
| INT-007 | Biofilm, unidentified composition, lamella | 8 | 3.02 |
| INT-008 | *H. sapiens* (fibroblast) lamella | 17 | 1.96 |
| INT-009 | *S. cerevisiae* (purified nuclei) lamella | 21 | 1.51 |
| INT-010 | *H. sapiens* lamella | 63 | 1.34 |
| INT-012 | *M. musculus* lamella | 40 | 1.63 |
| INT-013 | *H. sapiens* (RPE1) lamella | 17 | 1.57 |
| INT-014 | *H. sapiens* (Jurkat) lamella | 177 | 1.97 |
| INT-015 | *H. sapiens*, purified chromatin | 45 | 1.97 |
| INT-016 | *S. cerevisiae* lamella | 82 | 2.41 |
| REPORTS<sup>c</sup> | User-submitted reconstructed tomograms | 204 | - |

To ensure consistency across the training collection, we included only datasets for which either the raw fractionated tilt images or motion corrected tilt stacks were available, and reconstructed tomograms for these using an automated pipeline that employed AreTomo3^13,14^ for tilt series alignment and Warp^10,73^ for reconstruction and CTF determination. All tomograms were reconstructed at a common scale of 10 A per pixel as raw volumes and as even/odd volume splits. For convenience, we also used the data collection to train a general Noise2Noise-style^74^ network for denoising, using which we also generated a denoised version of every tomogram (see Methods).

### Generating training data for semantic segmentation of common cellular features

After assembling the training data collection, the goal was to generate reliable segmentation networks for a set of common cellular features. For macromolecular complexes - including ribosomes, microtubules, TRiC, nuclear pore complexes (NPCs), the vault complex, and actin filaments - we used a 3D U-Net architecture^75,76^. Specifically, we adopted the same architecture as used in MemBrain-seg^43^, changing only the training routine and the loss function depending on the abundance and morphology of the target feature. For organelle segmentations - including mitochondria, the cytoplasm, the nucleus, and the nuclear envelope - we used a 2D U-Net architecture instead, as these features are large in extent and can be reliably identified from individual slices, and because the 2D training routine is less computationally expensive than the 3D alternative.

Because we used experimental data, our dataset lacked any ground-truth labels. The main challenge in creating the segmentation networks was therefore the generation of the training labels. Since fully manual annotation is extremely time consuming, we used an iterative, network-assisted approach to generate training labels, a strategy which is similar to that employed in DeePiCt^38^. The procedure relied on Ais^39^ for manual, sparse annotation and for the training of preliminary 2D networks and on Pom^45,77^ to organise the training collection as a searchable database. This allowed us to browse the training collection, filter it by dataset of origin, and visualise the preliminary segmentation outputs alongside the corresponding tomograms, in order to identify errors in the outputs of the preliminary networks. By iteratively improving the labels, training, and testing against the large training collection, we achieved preliminary networks that produced qualitatively useful segmentations, as judged by visual inspection.

We would then explore the segmentation results again, this time in order to identify suitable locations to extract subtomograms for inclusion in the final 3D U-Net training data. In more detail, we marked manually sites either for the extraction of a ‘positive’ subtomogram or of a ‘negative’ subtomogram. The training labels for the negative subtomograms were all background, while the training labels for the positive subtomograms were generated using the preliminary 2D network in combination with post-processing steps that were specific to each feature of interest. These postprocessing steps are described in the online model library (mgflast.github.io/easymode/models). In short, they comprised a mixture of blurring, filament tracing or centroid detection (where relevant), thresholding and removal of small, disconnected regions, and depending on the feature of interest, downsampling to either 10, 20, 30, or 50 A /px. This last step was used to strike a balance between spatial detail, receptive field size, and processing speed. For example, the networks for fine-grained structures such as ribosomes, microtubules, and vault complexes were trained at 10 A /px, while networks for organelles like mitochondria and the cytoplasm were trained at a downsampled 50 A /px.

After creating a curated set of annotated subtomograms for training, we trained the 3D U-Net on this selection of labelled positive and negative subtomograms. To augment the training data, we applied standard transformations including rotations around the Z axis, rotations of up to 15 degrees around the X or Y axis, flips along the X or Z axis, contrast and brightness jitter, random rescaling (between 90% and 110%), and a mixup-style^78^ augmentation in which a randomly selected negative subtomogram is mixed with a positive training sample. In addition, each time a subtomogram is loaded during training, its density was sampled as a random linear combination of two of the four tomogram variants (even, odd, raw, and denoised), exposing the network to a range of different noise levels.

### Validation of the network outputs

Within cryo-ET, segmentations are most commonly used either for particle picking or to filter particle sets to specific subtomogram regions. To validate our networks, we therefore adopted two validation strategies that reflect how segmentation results are typically used in practice in cryo-ET workflows.

For features such as ribosomes and microtubules, which have a well-defined structure, we assessed network performance via downstream STA results. Specifically, we used the output of the segmentation networks to generate particle coordinate lists, which we then used for STA in the Warp/Relion5/M pipeline^10,11,17^. The rationale here was that, if applied to previously unseen data, the network produces coordinate sets that lead to high-resolution averages of the intended structure, then the network must be detecting the target feature with a reasonable degree of accuracy. A downside to this approach is that it does not quantify detection completeness, nor an absence of false positives; however, within common cryo-ET workflows, successful subtomogram averaging is one of the best possible ways to approximate some ground truth about the identity of a particle.

For features that cannot be meaningfully averaged, such as organelles and granules, we instead evaluated network performance using standard segmentation metrics (precision, recall, and Dice score).

In all cases, networks were tested on data that was either not included in the training collection at all, or on a dataset which we removed from the training collection for the sake of the test, or in some cases on a validation split of a dataset from which some other tomograms were included in the training collection.

#### In situ structure determination of protein complexes by subtomogram averaging

Subtomogram averaging can be used to resolve the three-dimensional structure of a macromolecular complex by averaging many instances extracted from tomograms, and is one of the principal applications of cryo-ET. The quality of the resulting reconstructions depends on the target complex being sufficiently abundant and on the accuracy with which individual particles are identified and localized. We used a segmentation-based particle-picking approach to perform STA of ribosomes, microtubules, TRiC, nuclear pore complexes (NPCs), vault complexes, and actin filaments, using easymode for segmentation and particle picking, and the Warp/RELION5/M pipeline for STA^10,11,17^.

In practical terms, segmentation-based picking for STA works as follows: one or multiple pretrained models are applied to segment a tomogram or dataset, producing segmentation volumes with the same dimensions as the input in which voxel values indicate the presence of each feature. These segmentation volumes are well suited for 3D visualization (**Fig. 1A-D**), which can help users choose appropriate picking parameters, such as the threshold, particle spacing, and picking mode. To derive particle lists from the segmentations, a number of picking modes are available, which are suited to different geometries: placing coordinates at the centroids of isolated particles (e.g. NPCs), at distance-transform local maxima (for connected or densely clustered particles, e.g. ribosomes), and filament tracing (for microtubles, actin, etc.). Some of these modes can also yield priors on the particle orientation (see Methods, **Extended Data Fig. 1**).

**Figure 1:**
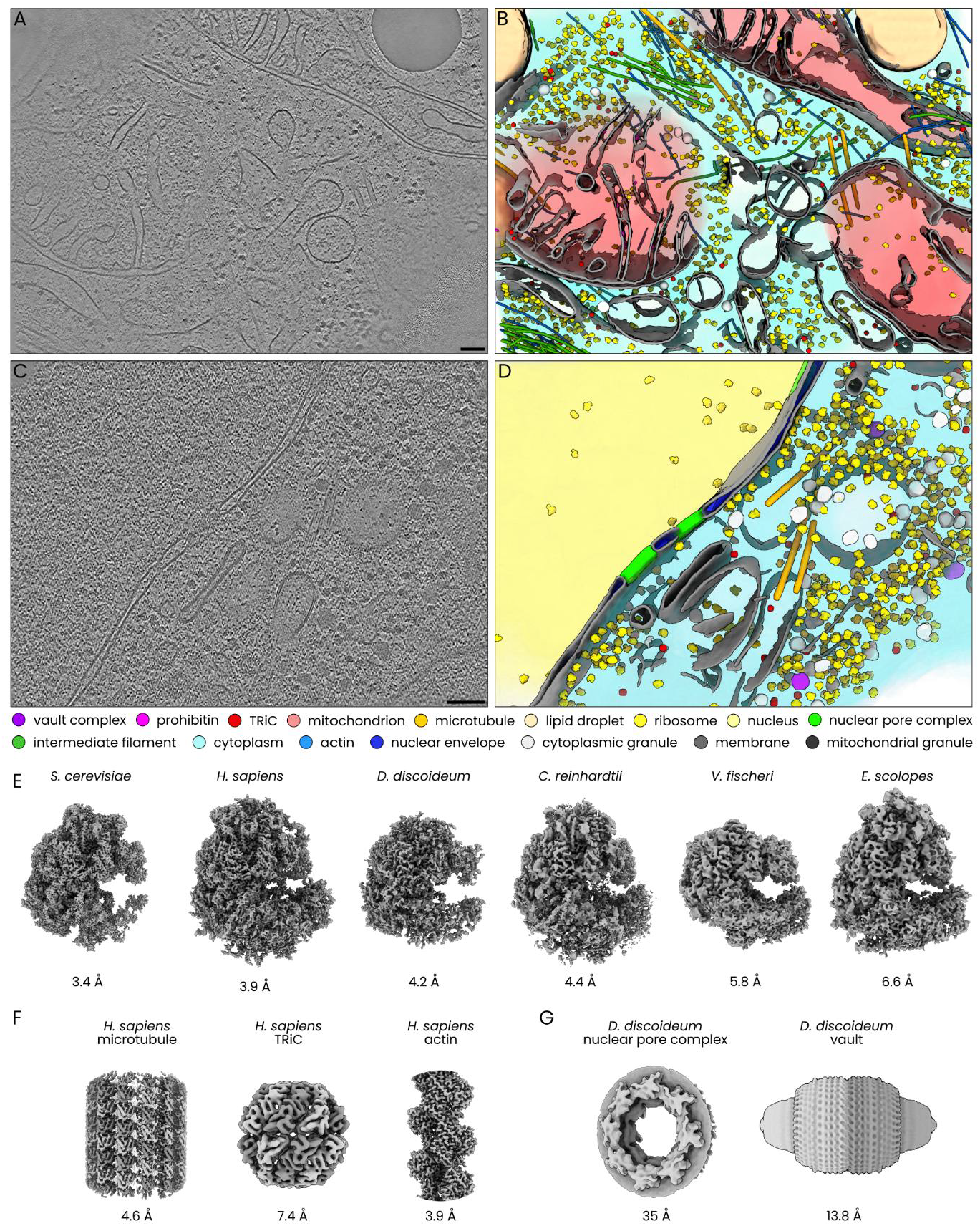
Validation of easymode networks by segmentation-based particle picking and subtomogram averaging. **A)** Tomographic slice of a *H. sapiens* (HeLa cell) lamella (EMPIAR-11561)^***59***^. Scale bar is 100 nm. **B)** 3D visualization of the *H. sapiens* tomogram, segmented using easymode (see legend below). **C)** Tomographic slice of a *D. discoideum* lamella (EMPIAR-11943^***58***^) cell. Scale bar is 100 nm. **D)** 3D visualization of the *D. discoideum* tomogram, segmented using easymode. The legend below the panel indicates the identity of the 16 different features. **E)** Subtomogram averages of ribosomes from six different species, picked using the same general ribosome segmentation network. FSC plots for these maps can be found in **Extended Data Fig. 2. F)** Subtomogram averages of *H. sapiens* microtubule, TRiC, and actin filaments. **G)** Subtomogram averages of *D. discoideum* nuclear pore complex and vault. FSC plots for the maps in panels F and G can be found in **Extended Data Fig. 3**.

Ribosomes are among the most common macromolecular complexes in cellular tomograms across species, and beyond being a structural target in their own right, their detection can also be useful as a measure of data quality and as a source of fiducial particles for improving tilt series alignment and CTF estimation^11,12^. Accordingly, the ribosome network is the most exhaustively tested of the easymode networks. Importantly, we performed STA without the use of 3D classification, so that the averaged maps reflect the full particle set produced by segmentation-based picking rather than a curated subset (**Fig. 1E**). Ordered from highest to lowest resolution, we tested the network in *S. cerevisiae* (3.4 A), *H. sapiens* (3.9 A)^25^, *D. discoideum* (4.2 A)^65^, *C. reinhardtii* (4.4 A)^42^, *V. fischeri* (5.8 A)^71^, and *E. scolopes* (6.6 A)^71^, reaching subnanometer resolution in all cases (**Extended Data Fig. 2**).

Next, for validation of the microtubule, TRiC, and actin segmentation networks, we used a variety of datasets acquired on *H. sapiens* samples. For microtubules we obtained a reconstruction at 4.6 A from a set of 621 tomograms collected on FIB-milled H. sapiens (HeLa) cells (**Fig. 1F**). The processing involved an initial per-filament averaging step to determine filament polarity (see methods). In order to ensure that the segmentation output for adjacent microtubules does not overlap, the postprocessing for the microtubule training data included a step to set the diameter of the tube label to 160 A, covering approximately the lumen of the microtubules only, which facilitated the tracing of individual filaments even when microtubules formed bundles.

For TRiC, we used a dataset of 360 high-magnification tomograms collected on *H. sapiens* (HEK293T) cells (EMPIAR-11538^25^), which was previously used to determine the *in situ* structure of TRiC^79^. Using the easymode model to segment TRiC, picking resulted in a set of 3202 particles (using data from the untreated condition only). Because TRiC occurs in a number of different conformations, STA of this complex needed 3D classification to separate the TRiC complex in the open and closed conformations^79^. We used one round of 3D classification into 4 classes to separate the particle set, which yielded a class with 620 particles that resembled the open conformation (as EMD-18922), two classes with 943 and 52 particles both resembling the closed conformation (as EMD-18927), and a final class of 1587 particles of which the average did not appear to correspond to the TRiC complex, although visual inspection of individual particles in this class suggested that a substantial fraction (105 out of the first 200 inspected) of these particles were genuine TRiC complexes that presumably did not get aligned correctly (**Extended Data Fig. 4-5**). Further refinement of the 943 particles in the closed state with D8 symmetry using RELION5 and M resulted in a map at 7.4 A resolution (**Fig. 1F**).

For actin, we imaged *H. sapiens* (fibroblast) ‘ghost cells’: cells treated with detergent to remove the membrane and soluble fractions, preserving the cytoskeletal network. The resulting samples allow the reconstruction of high-quality tomograms, suitable for STA of actin. Using a set of 24 tomograms, we segmented actin with easymode and traced filaments in the resulting segmentation, placing coordinates along them at a spacing of 150 A. This resulted in a selection of 29,000 particles, which averaged to a map at 3.9 A resolution (**Fig. 1F**) without the use of classification.

For STA of NPCs, we used the data from EMPIAR-11943 (**Fig. 1C-D**), comprising 130 tilt series of FIB-milled *D. discoideum* lamellae originally to study the dilation state of NPCs in different osmotic stress conditions^58^. Using the easymode NPC network we segmented all tomograms and picked particles, resulting in 174 coordinates without priors on the orientation. After an initial 3D refinement in RELION5^17^ to 55 A resolution, the particles were globally aligned to the reference, but without respect to the distinct cytoplasmic and nuclear sides of the complex. Using the easymode models for cytoplasm and nucleus segmentation to annotate the nuclear and cytoplasmic volumes, we then adjusted the orientation of each particle (**Extended Data Fig. 6**). A second 3D refinement then yielded a map of the NPC at 35 A resolution using C8 symmetry (**Fig. 1G**).

For the vault complex we used a dataset of 152 tomograms of FIB-milled *D. discoideum* lamellae (EMPIAR-11845^58^). To ensure an unbiased test, we excluded annotations in this dataset from the data used to train the vault network. Segmentation-based picking using the easymode vault yielded 393 particles, which averaged to a resolution of 13.8 A using D39 symmetry (**Fig. 1G**).

Taken together, these results demonstrate that these networks, in combination with segmentation-based picking strategies, are sufficiently accurate to support *in situ* structure determination by STA for this set of macromolecular targets, without requiring any per-dataset training or finetuning. Beyond STA of this handful of structures, the automated detection of these complexes – which are common to many (eukaryotic) species and often among the most abundant complexes in cellular tomograms – also enables other applications. This includes their use for data quality assessment, and as fiducials for improving tilt series alignment^11,12^. In combination with networks for the segmentation of organelles and other pleomorphic cellular features, the output of these networks can be used to visualize and build models of the cellular architecture observed in tomograms.

#### Validation metrics for organelle and utility segmentations

Organelle masks are commonly used in cryo-ET workflows to associate particles with their local biological context. This can be useful to filter particle selections — for example, to retain only template matching hits that fall within the expected cellular compartment — or to classify particles based on their subcellular localisation, as demonstrated effectively for nucleosomes within the nucleus^42^ or for ATP synthase, mitochondrial ribosomes, and heat shock proteins within mitochondria^41,80^. Generating the required masks by manual annotation can be very time consuming and impractical when datasets comprise hundreds or thousands of tomograms.

We therefore trained networks to segment a set of prevalent and important organelles: mitochondria, the cytoplasm, the nucleus, and the nuclear envelope. We also trained a network for cytoplasmic granules, which turned out to be a prevalent feature across many datasets in the training collection. Finally, we trained a network to detect the void — the region outside the biological sample — which effectively segments the sample boundary. For the organelle and void networks, we used 2D U-Net architectures operating at downsampled pixel sizes of 30 or 50 A /px, as these features are large in extent and can be reliably identified from individual slices. For the cytoplasmic granule network, we used the same 3D U-Net architecture as for the complexes described earlier, operating at a downsampled 20 A /px.

We then validated these networks by comparing their output to manually annotated ground-truth masks on data not used during training, spanning multiple species including *E. scolopes, S. cerevisiae, C. reinhardtii, H. sapiens*, and *D. discoideum*, training a separate instance of the network for each test with training data from the test dataset excluded (**Fig. 2A**).

**Figure 2:**
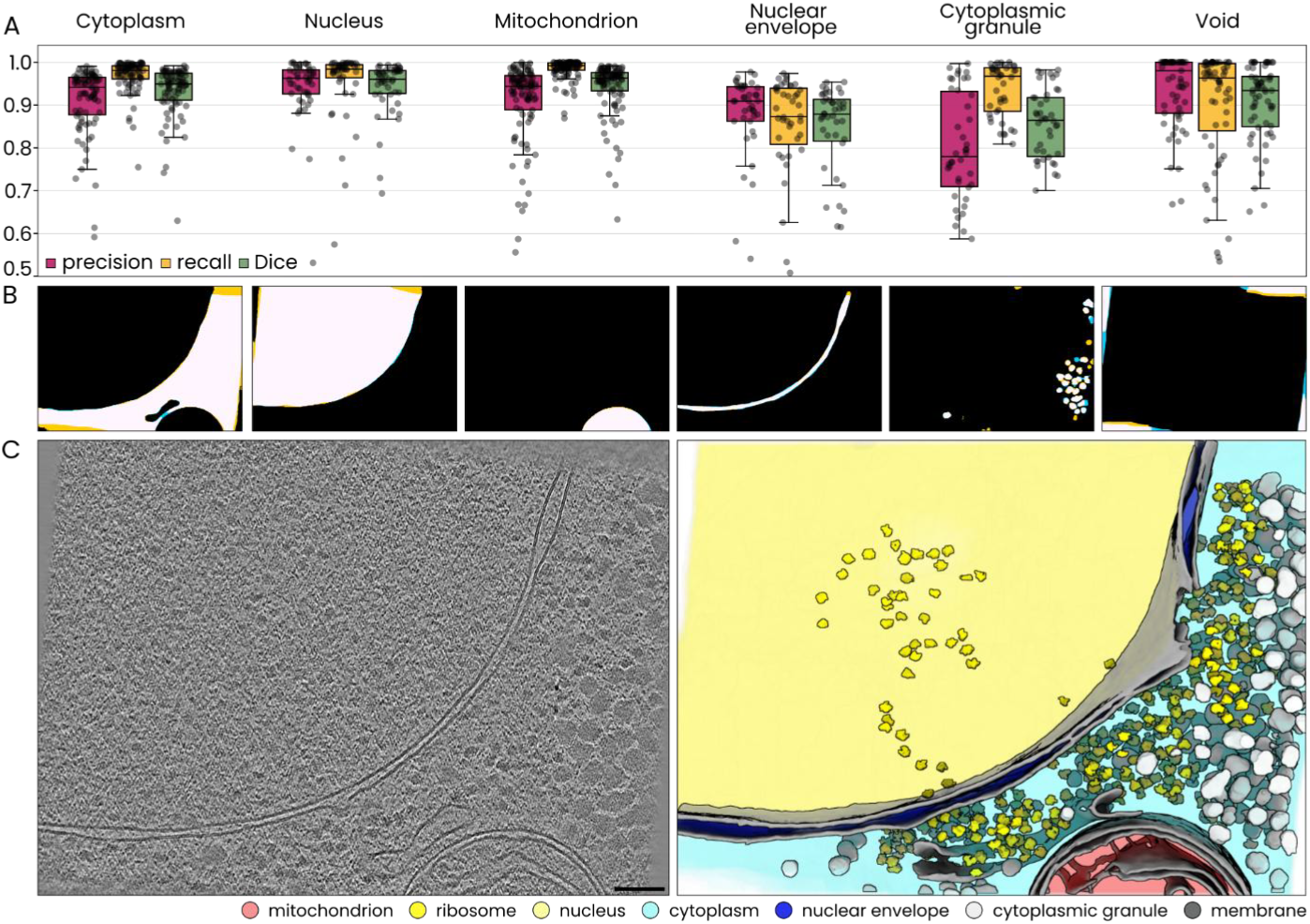
Validation of organelles, cytoplasmic granules, and void segmentation networks. **A)** Precision, recall, and Dice scores for six networks, each tested on data from at least four separate datasets and species. Boxes span the range from quartile 1 to quartile 3, with the median value marked. Whiskers extend to the most extreme value within 1.5x the interquartile range. All individual data points are shown. **B**) Examples of the network output (yellow) and ground truth masks (blue) for the feature labelled above. The white area is where the network output and ground truth labels overlap. **C)** The tomogram (*D. discoideum*, EMPIAR-11943^***58***^) corresponding to the example labels in B, showing the raw tomographic slice (left) alongside a 3D visualization of easymode segmentations of nucleus, cytoplasm, nuclear envelope, mitochondrion, cytoplasmic granules, ribosomes and membranes.

The cytoplasm, nucleus, and mitochondrion networks achieved mean Dice scores above 0.92 (n = 87, 47, and 103 tomograms, respectively, measured on central slices), precision above 0.90, and recall above 0.95, with particularly high recall for mitochondria (0.985 ± 0.023). The void network achieved a Dice of 0.890 ± 0.144 (n = 63), with more variable recall, reflecting in part the inherent ambiguity of the void boundary, where the transition between sample and non-sample is not sharply defined and differences in where this boundary is placed lead to lower overlap scores, particularly in slices with large void regions. Scores were lower for the nuclear envelope (Dice 0.823 ± 0.166, n = 43) and the cytoplasmic granules (Dice 0.784 ± 0.174, n = 42), both features where the ratio of edge to mask area is high, and where mask boundaries are less sharply defined (particularly for cytoplasmic granules), leading to larger overlap errors. Importantly, in all cases, errors were more common closer to the edges of the field of view, where the available spatial context is limited compared to more central regions.

To be noted is that the threshold used to binarise the continuous network output allows for some control over the segmentation boundary to suit a given downstream task; values reported here used a default threshold of 0.5, half-way between background and foreground, and lowering or raising this threshold shifts the balance between recall and precision (**Extended Data Fig. 7**). Example outputs for a single tomographic slice from a representative *D. discoideum* tomogram are shown alongside the corresponding tomographic slice and a 3D visualisation of the combined segmentations (**Fig. 2B–C**).

### Easymode pretrained general networks support workflows for in situ structure determination

Beyond quantitative validation, our primary goal is for easymode to be practically useful in cryo-ET workflows, including for *in situ* structure determination by STA. We present three exemplary cases in which easymode segmentations are used to improve commonly employed strategies: first, using organelle segmentations to filter particle sets from template matching; second, using segmentations to measure spatial relationships between particles and other cellular features and to filter particles accordingly; and third, using segmentation to define a picking strategy for a highly specific target, based on biological and geometric priors.

#### Filtering heterogenous particle sets based on cellular context

Template matching is a widely used strategy for particle picking in cryo-ET, in which a known reference structure is systematically cross-correlated against the tomographic volume to identify instances of the target complex^31–33^. The resulting correlation scores are then thresholded to yield a set of candidate particle coordinates and orientations. While template matching can be highly effective, the resulting particle sets are often heterogeneous: large, dark features, such as ice contamination, or membrane fragments, or structurally similar complexes frequently produce high correlation scores and are included among the hits. This problem can be addressed by 3D classification, but a more direct approach is to restrict picks to biologically plausible locations using organelle segmentation masks — for example, retaining only hits that fall within the nucleus when picking nucleosomes^42^. This approach requires segmentation of the relevant compartments, which in published work to date has often meant manual annotation^41,42^, a step that scales poorly with dataset size.

To demonstrate the use of easymode to this filtering step, we used a dataset of 239 tomograms from *H. sapiens* (HeLa) cells in which mitochondria are enriched in chaperones due to folding stress (EMPIAR-13145)^41^. Following the original approach of Ehses et al.^41^, we used PyTOM^33^ to perform template matching across all tomograms, which, with default particle extraction settings, yielded approximately 22,200 candidate particles. We then used easymode to segment mitochondria, membranes, ice particles, and ribosomes, and used these to define inclusion criteria to filter the particle set. We constrained the picks to the interior of mitochondria, and excluded regions within 50 A of any membrane, ribosome, or ice particle segmentation. These additional exclusions were intended to remove the small fraction of false positives arising from dense or high-contrast features within mitochondria. Filtering the particle set using this mask reduced it from 22,200 to approximately 3,500 particles (**Fig. 3C**).

**Figure 3:**
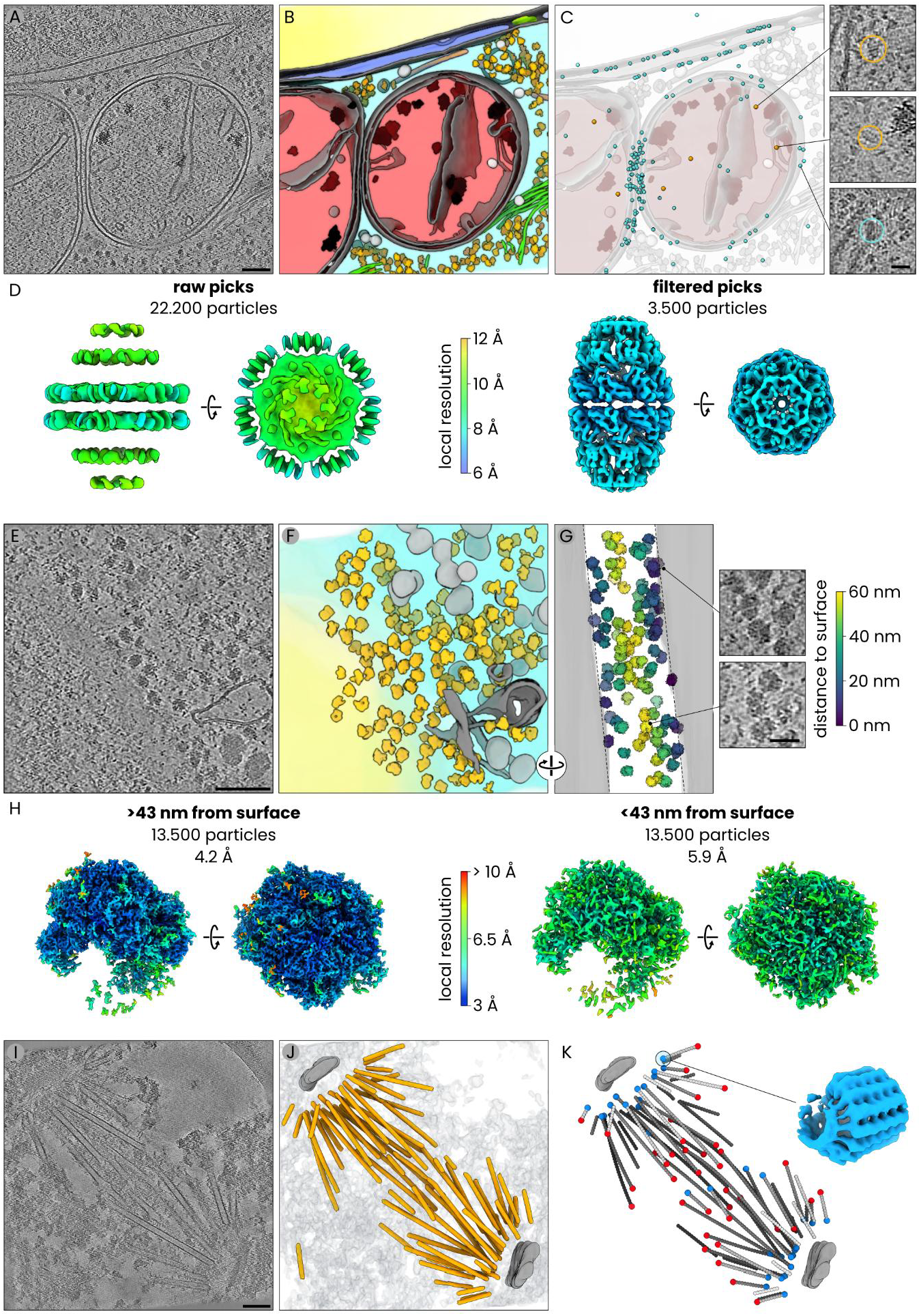
Pretrained segmentation networks support in situ structure determination workflows. **A)** Tomographic slice of a *H. sapiens* (HeLa) cell (EMPIAR-13145) under mitochondrial proteostatic stress^*41*^. Scale bar 100 nm. **B)** A 3D visualization of easymode network outputs, featuring ribosomes (orange), vimentin intermediate filaments (green), cytoplasmic granules (light gray), membranes (gray), mitochondrial granules (dark gray), microtubules (orange), cytoplasm (cyan), mitochondria (red), nucleus (yellow), nuclear envelope (blue), and a nuclear pore complex (lime). **C)** The location of template matching derived candidate Hsp60-Hsp10 complex coordinates, with false positives indicated in cyan and true positives in orange. The inset shows examples of two true Hsp60-Hsp10 particles and one false positive. **D)** A comparison of subtomogram averaging results using the full set of 22,200 particles (left) versus using the filtered set of 3,500 particles that were detected inside of mitochondria, away from membranes, ribosomes, or ice particles (right). The full particle set yielded an artefactual map at around 11 A, while the filtered set yielded a map of the Hsp60-Hsp10 complex at 8.0 A resolution. Maps are coloured by local resolution. **E)** Tomographic slice of a *D. discoideum* cell (EMPIAR-11899)^***65***^. Scale bar 100 nm. **F)** A 3D visualization of easymode network outputs, featuring ribosomes (orange), cytoplasmic granules (light gray), membranes (gray), nucleus (yellow), and cytoplasm (cyan). **G)** A side view of the same volumes, with ribosomes colour-coded by distance to the nearest lamella surface, which was determined using the easymode void segmentation output (gray). The inset shows examples of ribosomes located close to the lamella surface (top), or closer to the centre of the lamella (bottom). **H)** A comparison of subtomogram averages of the 50% of ribosomes located more than 43 nm away from the lamella surface versus the 50% found within 43 nm from the lamella surface, where damage due to cryoFIB milling affects the local structure. The colour bar indicates the local resolution. **I)** Tomographic slice of a mitotic spindle extracted from *S. uvarum*. Scale bar 100 nm. **J)** A 3D visualization of easymode microtubule segmentation, with segmentations of the spindle pole body (grey) and chromatin (transparent grey) generated in Ais added for context. **K)** A 3D visualization of the detected microtubule contours, with indicators for the plus-end (red) and minus-end (blue) detected via per-filament subtomogram averaging. A map of the γTuRC complex at 23.6 A resolution, determined via averaging of subtomograms extracted at microtubule minus ends, is shown in blue (see **Extended Data Fig. 11** for a map of the *S. cerevisiae* γTuRC complex fitted into the map).

To assess the effect of this filtering, we averaged both particle sets using identical RELION5/M procedures without any 3D classification (**Fig. 3D**). The unfiltered set of 22,200 particles yielded an artefactual map with a nominal resolution of ∼11 A but no genuine structural features, and was dominated by what appeared to be averaged membrane density. The filtered set of 3,500 particles refined to 8.0 A resolution (D7 symmetry) and clearly showed the secondary structure of the expected Hsp60-Hsp10 complex (**Extended Data Fig. 8**). In a second test, STA of nucleosomes also benefited from constraining template matching candidate particles to automatically segmented nucleus masks, revealing inner features of the nucleosome particles that are not visible when using the unfiltered particle set (**Extended Data Fig. 9**). These results demonstrate that pretrained segmentation networks for organelle and feature masking can replace manual annotation in template matching workflows, enabling automated contextual filtering of particle sets at dataset scale.

#### Filtering particles based on spatial proximity to sample boundaries

Beyond binary inclusion or exclusion from a compartment, segmentation masks can also be used to measure continuous spatial relationships between particles and other cellular or sample features, and to filter particle sets accordingly. We demonstrate this with an example in which the distance between each ribosome and the nearest lamella surface is used as a proxy for FIB milling-induced damage. Material close to the milled surfaces of a FIB-milled lamella is subject to ion beam damage, which degrades the local structure and reduces the achievable resolution in subtomogram averaging. This effect was characterised in multiple studies^4,65,81^, including in one by Tuijtel et al.^65^ whose dataset (EMPIAR-11899) we used for our test.

Using the easymode ribosome network to detect and pick particles and the void network to segment the lamella boundary, we identified approximately 27,000 ribosomes. After an initial refinement to ∼10 A resolution in RELION5, we measured the distance from each refined coordinate to the nearest void surface and split the set at the median value of 43 nm (comparable to ∼50 nm identified by Tuijtel et al.^65^) into two equally sized subsets: one comprising particles from the central region of the lamella, and one comprising particles closer to the milled surfaces. Simultaneous refinement of both particles sets in M with identical processing parameters for both subsets, we obtained maps at 4.2 A resolution for the central particles and 5.9 A for those nearer to the lamella surface (E**xtended Data Fig. 10**). This example illustrates how combining particle coordinates with continuous distance measurements derived from segmentation masks can be used to stratify particles by data quality, or more generally, how spatial relationships between detected features can inform downstream analysis decisions. While we demonstrate this here for distance to the sample boundary, the same processing steps, combined with different network outputs, are also applicable to other features — for example, to differentiate between membrane-bound or unbound complexes, or to measure particle location with respect to specific organelles (not shown).

#### Biological and geometric priors for particle picking

Targets of interest in cellular cryo-ET are often described in terms of their spatial and geometric relationship to other cellular features. When some of these features are readily detectable by segmentation, this information can be used not only to filter existing particle sets — as in the preceding two examples — but also to define the picking task itself as a combination of biological and geometric priors. We demonstrate this with the γ-tubulin ring complex (γTuRC), which caps spindle microtubules at their minus ends and serves as the site from which microtubules nucleate^82^. Microtubules can be readily and automatically detected using our pretrained network; detection of γTuRC, on the other hand, would require a dedicated approach such as template matching or training of a bespoke neural network. Its location can, however, be determined based on the geometry of the microtubule which it caps.

We demonstrate this approach using a dataset of 109 tomograms collected on purified mitotic spindles from the yeast *S. uvarum* (**Fig. 3I**). Using the easymode microtubule network we segmented and traced individual microtubules, picked particles along each filament, and averaged each filament individually in RELION5 to determine its polarity from the cross-section of the reconstruction (**Fig. 3J-K**). For each filament with an assigned polarity, we then traversed to its minus end and placed a candidate γTuRC coordinate at the terminal position, oriented along the tangent to the filament, with the filament polarity considered. To exclude filament ends arising from truncation or depolymerization at the sample surface rather than genuine microtubule origins, we discarded any coordinate within 250 A of the tomogram boundaries, detecting the top and bottom edges using the void network (example scripts for these steps are available online).

This picking routine yielded a set of 2,841 candidate γTuRC particles. One round of 3D classification using a large box size (768 A) and a half-microtubule reference then yielded a class of 701 particles that resembled γTuRC. Further refinement of this particle subset class resulted in a map at 23.6 A resolution (**Fig. 3K, Extended Data Fig. 11-12**). This example illustrates how pretrained segmentation networks can be used to formulate picking strategies based on biological and geometric context. Since these strategies require no reference map for the target of interest, they may offer a useful complementary or alternative picking strategy to direct detection methods such as template matching.

### Modelling cellular architecture with easymode: high-resolution in situ structure of IMPDH filaments

As a final example, we demonstrate the extensibility of the easymode network collection and its use in mapping cellular architecture by describing how easymode has contributed to our own cryo-ET study of enzymatic filaments in the purine synthesis pathway. While screening the training data collection and preparing annotations for the features described above, we sporadically observed filaments that did not belong to classical cytoskeletal filaments, but resembled filaments of inosine monophosphate dehydrogenase (IMPDH)^83,84^-the rate-limiting enzyme in de novo purine synthesis - in tomograms from seven separate datasets (human cultured Jurkat, RPE1^38^, HEK293T^25^, HeLa^41^, U2OS, and fibroblast cell lines, and mouse embryonic stem cells^60^). We knew about the features of these filaments because we were already collecting a large dataset of tomograms of cryoFIB-milled human (HeLa) cells treated with mycophenolic acid - an immunosuppressive compound known to induce IMPDH filament formation - in order to study the composition and architecture of the large IMPDH filament bundles known as RR-structures^85^ or cytoophidia^86,87^. In line with the goal of growing the easymode model library over time, we used the training collection to prepare a network for IMPDH segmentation following the same iterative procedure described above, and applied the resulting network to our dataset of 1,034 tomograms (which required around 90 minutes of processing on 8 RTX3070 GPUs). Sorting the dataset by the volume of IMPDH detected within each tomogram, we identified 19 tomograms containing IMPDH filaments, ranging from isolated single filaments to dense bundles comprising thousands of subunits (**Fig. 4A-B**). Converting segmentations into particle coordinates yielded just under 10,000 particles, which with D4 symmetry refined (RELION5/M) to 4.0 A resolution, without the use of 3D classification (**Fig. 4C, Extended Data Fig. 13**). The resulting map was consistent with a model of the open-state IMPDH1 octamer^84^ (**Extended Data Fig. 14**).

**Figure 4:**
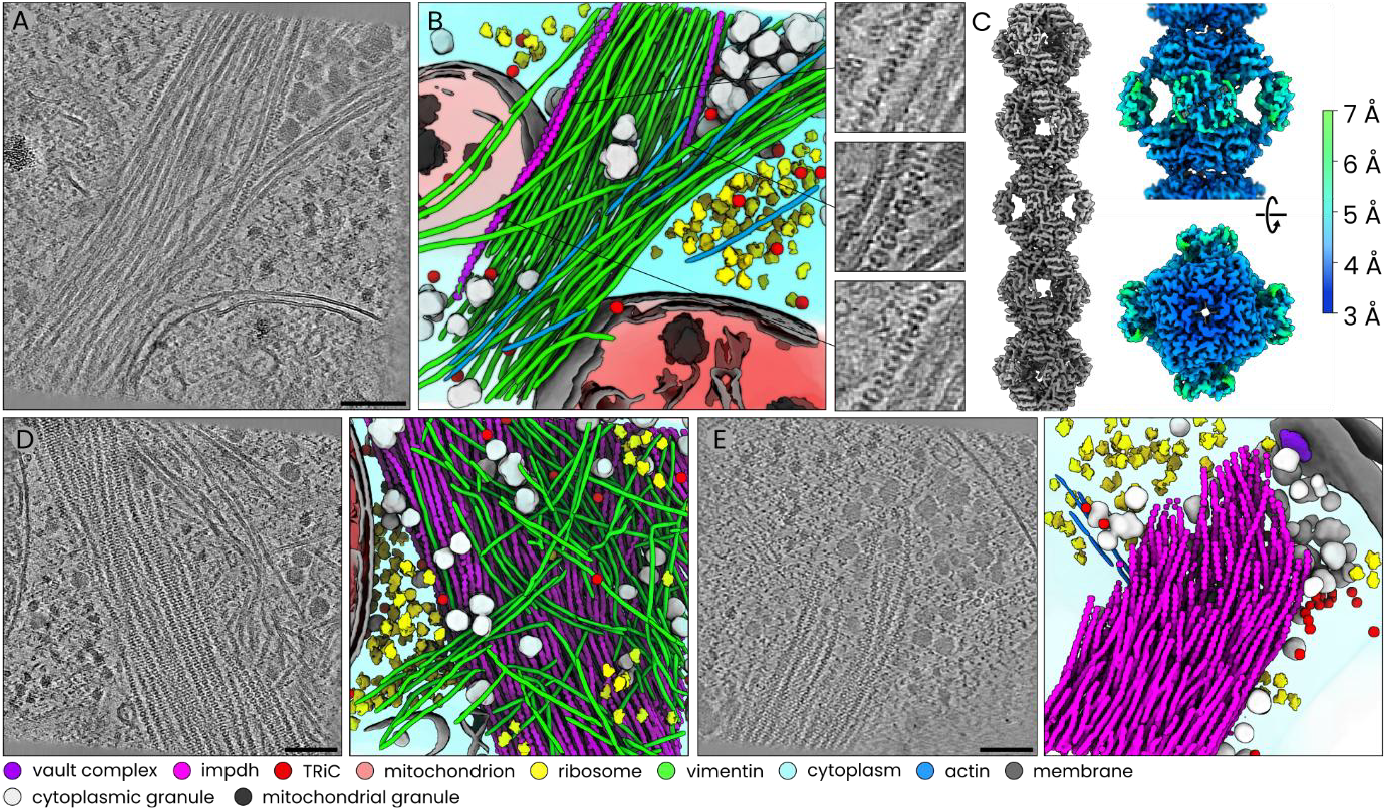
In situ structure of IMPDH filaments and mapping of their cellular context using easymode. **A)** Slice of a tomogram featuring three isolated IMPDH filaments (purple) in a bundle with intermediate filaments (green). **B)** A 3D visualization of easymode segmentation outputs, showing vimentin, IMPDH, ribosomes, TRiC, cytoplasmic granules, actin, membranes, mitochondria, mitochondrial granules, and the cytoplasm (legend at the bottom of the figure). **C)** In situ subtomogram average of IMPDH. Left: a composite map of five octameric repeats, spanning a little over one full turn of the helical assembly. Right: local resolution maps of the octameric repeat viewed from the side (top) and from within an octamer, looking at the C4 symmetric tetramer. **D)** An IMPDH filament bundle in association with many intermediate filaments. **E)** An IMPDH filament bundle in absence of intermediate filaments, surrounded by many cytoplasmic granules and TRiC complexes. All scale bars 100 nm.

To place these filaments in their cellular context, we then applied the broader set of easymode networks – for ribosomes, TRiC, actin, microtubules, vimentin intermediate filaments, membranes, cytoplasmic and mitochondrial granules, the vault complex, mitochondria, and the cytoplasm – to the same tomograms. Using these segmentations to map and visualise the cellular environment surrounding the IMPDH filaments, we find that individual filaments are frequently interspersed with a large number of intermediate filaments (**Fig. 4D**), the majority of which we putatively identify as vimentin filaments based on their diameter and appearance^88^. In tomograms containing large IMPDH bundles (**Fig. 4D-E**), these bundles appear to exclude other abundant cytoplasmic complexes such as ribosomes and TRiC. In some cases these bundles appear to consist of IMPDH filaments only, and are surrounded by many cytoplasmic granules (**Fig. 4E**).

Taken together, these results illustrate how general pretrained segmentation networks for cellular cryo-ET could be used by the growing community of structural cell biologists to help determine structures in their native context, explore datasets for features of interest, define picking strategies using biological and geometric priors, filter particle sets based on cellular context, and to visualize and model the major elements of cellular architecture based on tomogram volumes.

## Discussion

Here we present easymode, a collection of pretrained general segmentation networks for cellular cryo-ET that covers large cellular features, including cytoskeletal filaments, major organelles, prevalent complexes such as ribosomes, nuclear pore complexes, and less prevalent complexes like TRiC, the vault complex, IMPDH, and finally the detection of the sample boundaries. The networks can be applied directly to new data without requiring any per-dataset training or annotation: model weights are distributed via a HuggingFace server^89^, and users require no prior experience with machine learning. Together, the networks enable users to build maps of cellular architecture from tomographic volumes, to pick common complexes for subtomogram averaging, to filter particle sets based on cellular context, and to leverage biological and geometric priors for particle identification and STA.

### Design choices

Two design choices in this work warrant explanation. The first is that we used U-Net architectures^75,76^ for all networks in the collection. This was motivated by two observations: first is that U-Nets are well established and reliable for biomedical image segmentation, including in cryo-ET^37,38,43^. The second was that all of the top-performing submissions to the recent CZII ML particle picking challenge^90^ used U-Nets, confirming their effectiveness for this domain. In general, our main goal was to build an effective tool for cryo-ET workflows, which we see as a matter of translating established tools from machine learning into the domain of cryo-ET. Therefore we did not explore the potential of newer architectures for this purpose.

The second design choice is that all networks were trained exclusively on experimental data. Although simulated tilt series have proven valuable for training in several contexts^34,35,91–93^, we chose to use experimental data only for three reasons. First, the volume of publicly available cryo-ET datasets has grown substantially in recent years, and we considered there to be sufficient data available to support the training of general models. Second, training on real data eliminates the need to bridge the domain gap between simulated and experimental images. Third, some of the features we aimed to segment are difficult to simulate faithfully — for example, the complex membrane morphologies found in mitochondria. A consequence of training on real data is that all training labels were generated by manual annotation, which, for some features, is relatively straightforward (microtubules and mitochondria, for instance, are large and visually unambiguous) but for others, such as actin filaments viewed in cross-section, can be difficult. For these harder cases, an approach that combines real and simulated training data could improve detection quality^93^, and is something we aim to explore in future iterations of the networks.

### Applications

Cellular tomograms contain a wealth of structural information beyond the specific target of any given study, and many biological questions can be framed in terms of common features such as whether a complex localizes to the nucleus, the cytoplasm, to mitochondria, or some other organelle, and whether it associates with membranes, or perhaps the cytoskeleton. Making these features computationally accessible could therefore substantially improve our ability to leverage the detailed biological information offered by cryo-ET in order to study target complexes in their native environment. The examples discussed in this work illustrate several ways in which easymode can be used in practice.

The most direct application is particle picking of the common complexes covered by a pretrained network (**Fig. 1**). For these, segmentation followed by coordinate extraction provides a straightforward alternative to template matching (TM) or manual picking. Using this approach, we obtained subtomogram averages of ribosomes, microtubules, actin filaments, and TRiC with sub-nanometer resolution, in all cases for novel datasets to which the networks had not previously been exposed. Compared to TM, segmentation-based picking of these targets is fast and produces particle sets that may be refined without extensive 3D classification. The TRiC example illustrates the practical trade-offs between TM and segmentation-based picking most clearly. In their original work, Xing et al.^79^ used TM to pick 355,000 candidate particles, which after extensive 3D classification reduced to 4,054 and yielded a map at 7.1 A resolution. In our work, using the same dataset of 355 tomograms, our segmentation-based picking yielded ∼3,200 particles initially, which reduced to ∼943 in a single round of 3D classification, and yielded a map at 7.4 A resolution. Using TM settings comparable to the original, we measured that the TM-based search takes around 370 seconds per tomogram, compared to 16 seconds per tomogram for the segmentation (tested on the same hardware). This comparison illustrates that segmentation-based picking can be an efficient alternative to TM in cases where pretrained networks are available. A more detailed overview of processing speed with easymode can be found in the supplementary information (**Extended Data Fig. 15**). For the most common network (3D UNet, 10 A /px), which is used for most macromolecular complexes, including ribosomes, microtubules, and actin, segmentation of the IMPDH-filament dataset (**Fig. 4**) took around 50 seconds per volume per GPU (volume size 310 × 620×620). The networks operating at larger pixel sizes are much faster; for example, the mitochondrion and cytoplasm segmentation (2D UNet, 50 A /px) took around 2 seconds per volume per GPU for this dataset.

For targets not covered by a pretrained network, segmentation outputs can be used in combination with other picking approaches to filter particle sets. We demonstrated this by using organelle segmentations to constrain TM-derived candidate Hsp60-Hsp10 particles to the interior of mitochondria (**Fig. 3A-D**) and of nucleosomes to nuclei (**Extended Data Fig. 9**), which in both cases substantially improves the result of subtomogram averaging. These results demonstrate how easymode can be used to replace manual annotation of regions of interest with automated segmentation. Segmentation outputs can also be used to measure spatial relationships between particles and other cellular features: we showed this by stratifying ribosomes by their distance to the lamella surface in order to separate damaged particles from those particles better preserved in the lamella interior (**Fig. 3E-H**). This logic, of using segmentation masks to define biological and geometric priors on particle identity, can also be extended to define picking strategies for novel particles. This was illustrated in the γTuRC example (**Fig. 3I-K**), where we used microtubule segmentations to trace filaments, determine their polarity, and place candidate particles at microtubule minus ends, yielding a 23.6 A average of γTuRC without requiring a reference map for the target itself.

Finally, by combining the outputs of many different networks, easymode can be used to visualize and explore the cellular context surrounding targets of interest. In the example of IMPDH, we initially applied just the IMPDH network to a set of over 1000 tomograms, and used the outputs to identify a small subset of tomograms containing IMPHD. While the IMPDH network specifically enabled particle picking and subtomogram averaging to 4.0 A resolution (**Fig. 4**), we then also applied the broader set of easymode networks to these tomograms to map the surrounding environment. The results suggested that IMPDH filaments are often associated with intermediate filaments, while bundles of IMPDH appear to exclude ribosomes and TRiC from their interior. Whether these spatial associations reflect specific biological interactions is beyond the scope of this work, but the availability of pretrained networks makes it much easier to raise and investigate such questions. In our own work, we also find this kind of segmentation-guided exploration especially useful in combination with the dataset-scale visualization and curation tool Pom^77^. After producing segmentations with easymode, Pom can be used to summarize the results across a large dataset, generate 2D and 3D visualizations of tomogram contents, and then present these in a searchable browser-based interface. This combined usage is also demonstrated in the online documentation.

### Limitations of this work

Some limitations of this work should be noted. First, we emphasize that easymode is not a tool for picking arbitrary targets. For the specific complex of interest in any given project, a pretrained network will not always be available, and detection will require tools such as template matching^31–33^, embedding-based tools such as TomoTwin^34^, ProPicker^36^, or cryoSIAM^35^, or other dedicated approaches^39,94^. By providing reliable, general networks for the detection of prevalent complexes and organelles, easymode enabled automated mapping of the cellular environment and renders the biological context of cryo-ET datasets computationally accessible, facilitating users to leverage this context in their analyses. For example, it offers the ability to screen large datasets for specific features (e.g., the presence of actin filaments) before selecting a subset of tomograms for more computationally expensive processing; to average common complexes such as ribosomes as a rapid assessment of data quality or to inform improved tomogram reconstruction^11,12^; to filter particles sets to biologically plausible compartments; and to help represent datasets in a visual, searchable way^45^. Especially in combination with other tools for picking arbitrary targets – such as in the example of the Hsp60-Hsp10 complex – easymode thus supports workflows for *in situ* protein structure determination.

Second, we note that validation via subtomogram averaging is not a flawless way of validating network output. While it directly addresses one of the main applications of cryo-ET, it does not provide a quantitative assessment of detection accuracy. High-resolution STA only demonstrates that a sufficient number of true positives are present in the particle set, but it does not quantify the number of false negatives, nor guarantee the absence of false positives. Nevertheless, high-resolution averages can be obtained only if a substantial fraction of the input particles are correctly picked. During the 3D classification of the TRiC particle set, we noticed that classification does not specifically discard false particles only, but rather it discards particles which do not contribute to the given class averages. Inspecting the discarded particles in the original tomograms, we found that for more than half of the cases we would in fact have included them in the particle set with manual picking (**Extended Data Fig. 5**). This problem of differentiating between genuine particles, false positives, and misaligned true positives reflects a fundamental difficulty in experimental cryo-ET, where every individual particle is a noisy instance of some underlying structure that can only be approximated via averaging. This is why, in our validation, we decided to focus on demonstrating that the segmentation-derived particle sets can be used in practice to produce high-resolution subtomogram averages. When using any of the easymode networks, it is important to be aware of these limitations.

Third, while the scale and diversity of the training collection and the iterative routine of training and testing against this large data collection were designed to yield networks that generalize broadly, network performance will naturally vary across datasets. Some failure modes are feature-specific: the quality of actin segmentations, for example, appears to depend strongly on tomographic contrast, and labels for individual filaments are often discontinuous in thicker or lower-dose volumes. Moreover, filaments oriented along the Z axis are not detected at all. Another example, the nuclear envelope network currently fails to detect the NE when it lies approximately in-plane with the lamella and the bounding membranes are not directly visible in the missing wedge-affected reconstruction. In both these cases, these limitations are ultimately due to the fact that we were unable to confidently annotate such examples during label preparation. More generally, because all labels were generated by manual annotation, the subjectivity inherent in this process propagates into the resulting models. Where we are aware of specific flaws in any model’s output, we have listed these in the model library. In practice, the most effective way to assess whether these limitations affect a given use case is to simply inspect the segmentation outputs alongside the corresponding tomograms, for example by using ChimeraX^95^ or Pom^45,77^.

Finally, the training collection itself is also not free of bias: nearly half of the datasets are from human samples. In practice, however, comparing volumes from *H. sapiens, M. musculus*, and *D. discoideum* datasets, we find that the main sources of variety between datasets are the imaging parameters such as the pixel size, dose, and defocus, rather than variation between species (**Extended Data Fig. 16**). A potentially more important source of variation in the training data is the sample preparation method. CryoFIB milled lamellae, the periphery of whole cells, and purified preparations all tend to produce orientation-biased views of cellular content, with larger structures such as filaments mostly seen in side projections. The training collection comprises a majority of samples of these types, and a smaller number of samples prepared by plasma FIB-milling in combination with lift-out^66,96,97^ or by CEMOVIS^63,98^. These alternative preparation techniques can produce samples with less orientation bias and can be used to prepare samples from tissues^64,99^ or whole organisms^71,97^, which may present features not well represented in the current training collection. As the diversity of publicly available cryo-ET data grows, we will expand the training collection accordingly, and we invite users to submit challenging examples via the easymode reporting tool, with which they can voluntarily upload tomograms to our data server for incorporation into future training iterations.

### Outlook

The studies highlighted in the introduction (and many others we did not mention) illustrate how cryo-ET can be used to bridge high-resolution *in situ* structural biology with the larger scale of cell biology. As datasets grow to the scale of thousands of tomograms^19,42^, the potential to extend the use of cellular cryo-ET beyond individual complexes and towards systematic descriptions of cellular organization becomes increasingly tangible. Realizing this potential requires not only the collection of large datasets, but also the ability to comprehensively map the biological features present within them – and as data volumes continue to increase, this will depend increasingly on automated and general computational tools. The widespread adoption of pretrained general networks such as MemBrain^43,44^ for membrane segmentation demonstrates the practical impact that such tools can have. By extending this approach to a broader set of common cellular features, we hope that easymode will help researchers in the cryo-ET community make the biological contents of their data more readily accessible.

The software, network weights, and documentation are freely available via mgflast.github.io/easymode, and will be maintained and extended over the coming years.

## Methods

### Tomogram reconstruction

All tomograms were reconstructed from fractionated tilt images when available or otherwise from motion corrected tilt stacks using WarpTools^10,73^ and AreTomo3^13^ (version 2.1.10), called via the command ‘easymode reconstruct’ to automate the processing. All tomograms were reconstructed in three variants: two volume half splits, the raw tomogram, and a denoised variant (see below). For datasets with fractionated tilt images available we used frame-based splitting for the volume half splits. For datasets with post-motion correction tilt stacks we used tilt-based splitting.

### Denoising with a general noise2noise-style denoiser

All tomograms were denoised using the easymode general Noise2Noise-style^74^ denoiser, a 3D residual UNet (∼8M parameters) that is included in the software. This network was trained in two steps: first, subtomogram pairs were extracted from even/odd frame-split^16^ tomograms across all datasets in the training collection for which raw frames were available, sampling an equal number of training volumes from each dataset. The network was then trained on these pairs and the resulting model subsequently applied to all even/odd training pairs to produce denoised versions of these training subtomograms. We then trained a second instance of the network using raw/denoised subtomogram pairs. We note that training a Noise2Noise model in this way is not strictly statistically sound, and that it may always be better to train a bespoke network (with cryoCARE^16^, for example) on one’s own data to ensure optimal performance. However, a benefit of the ‘direct’ denoiser is that it can be applied to full tomograms directly, bypassing the need to generate or run inference on independent half splits. We find that it produces qualitatively useful denoised outputs, while also helping to save memory and computation time.

### Generating training labels

The training data collection was organized as a browser-based searchable database using Pom^77^, which allows for the creation of tomogram subsets based on user-specified tags. In this case we tagged tomograms by dataset of origin and species type. When starting out annotation for a new feature, we used the data browser to select tomograms containing this feature, opened these tomograms in Ais^39^, and manually annotated one or multiple 2D slices of the denoised version of the tomogram. We then placed boxes, marking square regions of interest for extraction for the initial 2D training datasets. We also included such training boxes in tomograms that did not contain the feature of interest, for use as counter-examples during training; these were often samples from tomograms that were guaranteed not to contain the feature of interest (for example, when annotating nuclei, any tomogram of a prokaryotic sample can be used as a counter-example). After annotation, we used Ais to extract these boxes and scale them to the desired pixel size (10, 20, 30, or 50 A /px, depending on the feature), with a final box size of 128 × 128 pixels. After training a network using this data, we applied the resulting network to a relevant subset of the entire training data collection and used Pom to summarise the results and to prepare 2D and 3D visualizations of the segmentation output. These were then also visible in the data browser alongside the corresponding tomograms. We used this view to identify erroneous network outputs, and generally used this data browser to sort all tomograms in the training collection by the volume fraction of the feature of interest detected within each tomogram, in order to uncover additional volumes for annotation. After multiple rounds of iterative annotation and training, we achieved networks that were either directly suitable for use (2D case) or that could be used to prepare the 3D training data (for most macromolecule segmenting networks). In the latter case, we explored the tomograms alongside 3D isosurface visualizations of network outputs and manually selected 3D subtomograms (160 × 160×160 voxels at 10, 20, or 30 A /px) for post-processing and 3D network training. Of 4121 tomograms in the easymode training data collection, a total of 1677 were (sparsely) annotated, while all were used to test networks against during the iterative process of improving the networks.

### Network architecture and training

We used U-Net architectures for both the 2D and 3D networks. The 2D network consisted of four encoder and four decoder blocks. Each encoder block contained three convolutional layers (3×3 kernels, ReLU activation, batch normalization), followed by 2×2 max pooling. Filter counts doubled at each level from 64 to 512, with a 1024-filter bottleneck. Dropout (25%) was applied at the bottleneck. Each decoder block used transposed convolution for upsampling, concatenation with the corresponding encoder feature maps, and two convolutional layers with the same kernel size, activation, and normalization. The output layer used a 1×1 convolution with sigmoid activation. Input patches were 128×128 pixels. The 3D network mostly followed the residual U-Net architecture used in MemBrain-seg^43^. The encoder consisted of five residual blocks (two convolutional layers each, 3×3×3 kernels, group normalization with 8 groups, ReLU activation), with strided convolution for downsampling. Filter counts increased from 32 to 512 across encoder levels, with a 1024-filter bottleneck. The decoder mirrored the encoder with five residual blocks and transposed convolution for upsampling. A 1×1×1 convolution with sigmoid activation produced the output. Input subvolumes were 160×160×160 voxels. All networks were trained with the Adam optimizer and a combined binary cross-entropy and Dice loss, with weights 0.2 and 0.8, respectively. A loss-ignore mask excluded a 16-pixel border for 2D training patches and a 32-voxel border for 3D subvolumes. Training was performed on 4 or 8 NVIDIA A100 GPUs with a batch size of 8–16 per GPU (2D) or 2 per GPU (3D). We note that we employed a single training strategy across all features, varying only the network dimensionality, pixel size, and post-processing of training labels. This reflects a deliberate focus on breadth of coverage rather than per-feature optimization. Feature-specific tuning of architecture, loss function, or training schedule could likely yield further improvements on individual segmentation tasks. All code is publicly available; most of the training data will be made publicly available upon publication, and is available upon request in the interim.

### Sample preparation and data acquisition for the IMPDH and actin datasets

HeLa cells were seeded on plasma treated R2/1 Quantifoil 200 mesh gold grids in DMEM supplemented with 10% fetal bovine serum and 100 U/mL penicillin and streptomycin, under normal culture conditions (5% CO_2_, 37 °C). After 24 hours we added mycophenolic acid to a final concentration of 100 µM and incubated the culture for another 16 hours before plunging grids into liquid ethane (Leica EM GP, 90% humidity, 22 °C chamber, 10 – 15 second blot). Grids were then clipped and loaded into an Aquilos 2 cryoFIB for lamella preparation using automated milling in AutoTEM^100^. Tilt series were acquired on a Titan Krios 300 keV microscope using a Falcon4i and SelectrisX energy filter at a pixel size of 1.514 Å and a total dose of 150 electrons/Å^2^ over a dose-symmetric tilt range from −45° to +45° in 3° increments, centred on the milling angle of 10°, and with a nominal defocus of 2.0 - 5.0 µm.

For the ghost cells, Quantifoil R1.2/1.3 gold 300 mesh grids were plasma treated then coated with fibronectin (50 ug/mL in PBS) for 45 minutes, then washed with PBS before seeding human skin fibroblasts (immortalised with SV40LT and TERT) and incubating grids for 18 hours. Grids were then washed with PBS and treated with permeabilization buffer (50 mM MOPS pH 7.0, 10 mM MgCl_2_, 0.15% Triton X-100 in PBS with protease inhibitors) for 3 minutes. Benzonase (Merck Millipore) was then added to a final concentration of 2.5 U/uL and the grids were incubated for 30 minutes at room temperature. After washing in PBS, grids were plunged as described above (2 second blot). Tilt-series were acquired on the same microscope as above at a pixel size of 1.96 A using a dose-symmetric acquisition scheme between −60° and +60°, 3° increments, a total dose of 100 electrons/Å^2^, and a nominal defocus of 2.0 – 5.0 µm.

### Subtomogram averaging

Particle picking was performed using the filament tracing and globular picking methods available in easymode/Ais^39^. For each target, two main parameters can be adjusted: (1) `size`, the minimum volume (in Å^3^) of connected components in the thresholded segmentation, and (2) `spacing`, the minimum distance between coordinates. For tracing filaments, the relevant parameters are slightly different; a `length` threshold for the minimum contour length of a traced filament is used instead of size, and coordinates are placed along filament contours at defined `spacing`. A third parameter, the threshold, was mostly kept at the default value of 0.5, but in practice can be adjusted to overpick or instead pick more selectively.

All subtomogram averaging was done using the Warp/RELION5/M pipeline^10,11,17^, with RELION5 used for initial pose estimation and M for refinement. The core of this workflow is described in detail online^73^. For initial averaging in RELION5, particles were extracted using WarpTools command `‘ts_export_particles’` using a pixel size of 4 – 5 Å /px and the output starfile from `‘easymode pick’` as the input coordinates. Where possible, we refrained from using 3D classification or other methods of filtering particle sets. For microtubules, classification involved a step of manually determining filament polarity and protofilament number based on the cross-sections of per-filament averages. Filaments for which polarity was not clearly identifiable were excluded from the final particle set. For TRiC, one round of 3D classification in RELION5^17^ was used to isolate the subset of closed-state particles. For NPCs, we indirectly classified particles during the orientation-correction step by measuring the average cytoplasm and nucleus segmentation values in front of and behind each initial NPC position-orientation estimate, and discarding particles when these values gave no consensus on the pore orientation. Step-by-step tutorials for the STA procedures described in this paper are available as part of the online documentation at mgflast.github.io/easymode

During refinement in M^11^ (used in all cases except for NPCs and γTuRC) we initially refined only the particle poses, then added image warp (up to 4 × 4) as resolution improved and CTF refinement once maps reached sub-7 Å resolution. In some cases, continued refinement in M would result in maps with a nominally improved resolution but poorer quality (increased noise, overly sharp features). Whenever we saw such signs of over-refinement, we reverted to data from earlier iterations and stopped refinement there.

### Segmentation and visualization

All segmentation was performed using easymode pretrained general networks, except for the spindle pole body and chromatin segmentations in Figure 3 which were prepared with Ais^39^. Visualizations in the main figures were created in ChimeraX^95^ and with the ArtiaX^101^ plugin (Fig. 3C, G, K). For 3D visualization of segmentation network outputs, post-processing steps included gaussian blurring along the Z-axis (for volume visualization of cytoplasm, nucleus, and mitochondrion segmentations) and use of the ‘remove dust’ function to remove small disconnected nonzero regions that were significantly smaller than the corresponding feature, analogous to the ‘size’ parameter used in picking.

### Particle contextualization

Segmentation-based particle contextualization was performed using Pom, with the function `‘pom contextualize’`. This command takes a particle starfile and any number of ‘samplers’ as the input. For example, for the Hsp60-Hsp10 filtering, one sampler was ‘v:mitochondrion:500’, which defines a volumetric measurement of the average mitochondrion segmentation value within a sphere of 500 A radius centred on each particle. Another sampler in this case was ‘r:membrane:0.5’, which defines a measurement of the distance to the nearest non-zero region in the membrane segmentation thresholded at a value of 0.5. This command writes the resulting values to new columns in the starfile, based on which the particle set can then be filtered. In the same Hsp60-Hsp10 example we discarded all particles for which the mitochondrion volume sampler returned less than 0.7 (1.0 being the maximum value) or for which the membrane sampler returned less than 50.0 Å. The contextualization tool is documented along with the main programme on mgflast.github.io/easymode.

## Acknowledgements

We would like to thank the authors of many private and public data contributions, including Tom Dendooven, A lia dos Santos, Piotr Kolata, Alexander Scrutton, Forson Gao, Cong Yu, Paula Paredes Vergara, Kashish Singh, Eric Wang, Andriko von Ku gelgen, Maite Freire Delgado, Mike Sleutel, David Barford, Katrina Gundlach, Oda Schiøtz, Ariane Briegel, Ju rgen Plitzko, Sebastian Tacke, Elisa Lisicki, Tatjana Taubitz, and Stefan Raunser, as well as the authors and depositors of the various EMPIAR and CDP datasets used. We also thank Shaoxia Chen, Giuseppe Cannone, Grigory Sharov, Anna Yeates, Bilal Ahsan, and Haaris Sadari of the LMB electron microscopy facility for help with cryoFIB milling and cryo-ET data acquisition, and Jake Grimmett, Toby Darling, and Ivan Clayson of the LMB scientific computing team for help with data processing. We acknowledge Diamond for access and support at the Aquilos-II and Krios of the UK national electron Bio-Imaging Centre (eBIC). For the purpose of open access, the MRC Laboratory of Molecular Biology has applied a CC BY public copyright licence to any Author Accepted Manuscript version arising. M.A. was funded by the UKRI Medical Research Council (MC_UP_1201/29), M.S. was funded by an EMBO postdoctoral fellowship (ALTF-721-2024).

## Data availability

Subtomogram averaging results for the following strucutres are available via the Electron Microscopy Data Bank (EMDB)^102^: IMPDH (EMD-58147), γTuRC (EMD-58163), Hsp60-Hsp10 (EMD-58187), vault complex (EMD-58166), NPC (EMD-58184), actin (EMD-58186), TRiC (EMD-58185), microtubule (EMD-58189), *S. cerevisiae* ribosome (EMD-58196), *H. sapiens* ribosome (EMD-58168), *D. discoideum* ribosome central (EMD-58194) and lamella surface (EMD-58195) subsets, *V. fischeri* ribosome (EMD-58191), and *E. scolopes* ribosome (EMD-58190).

## Author contributions

M.S. conceived the project; M.S. designed the analysis strategies supported by A.B and M.A.; M.S. prepared samples for and collected the MPA-treated HeLa dataset; T.H. prepared samples for and collected the ghost cells tomograms; M.S. developed the software, created annotations, analysed the data, performed subtomogram averaging and validation; M.A. supervised the study; M.S. wrote the manuscript with the contribution of all the authors.

## Supplementary Information

**Supplementary Figure 1:**
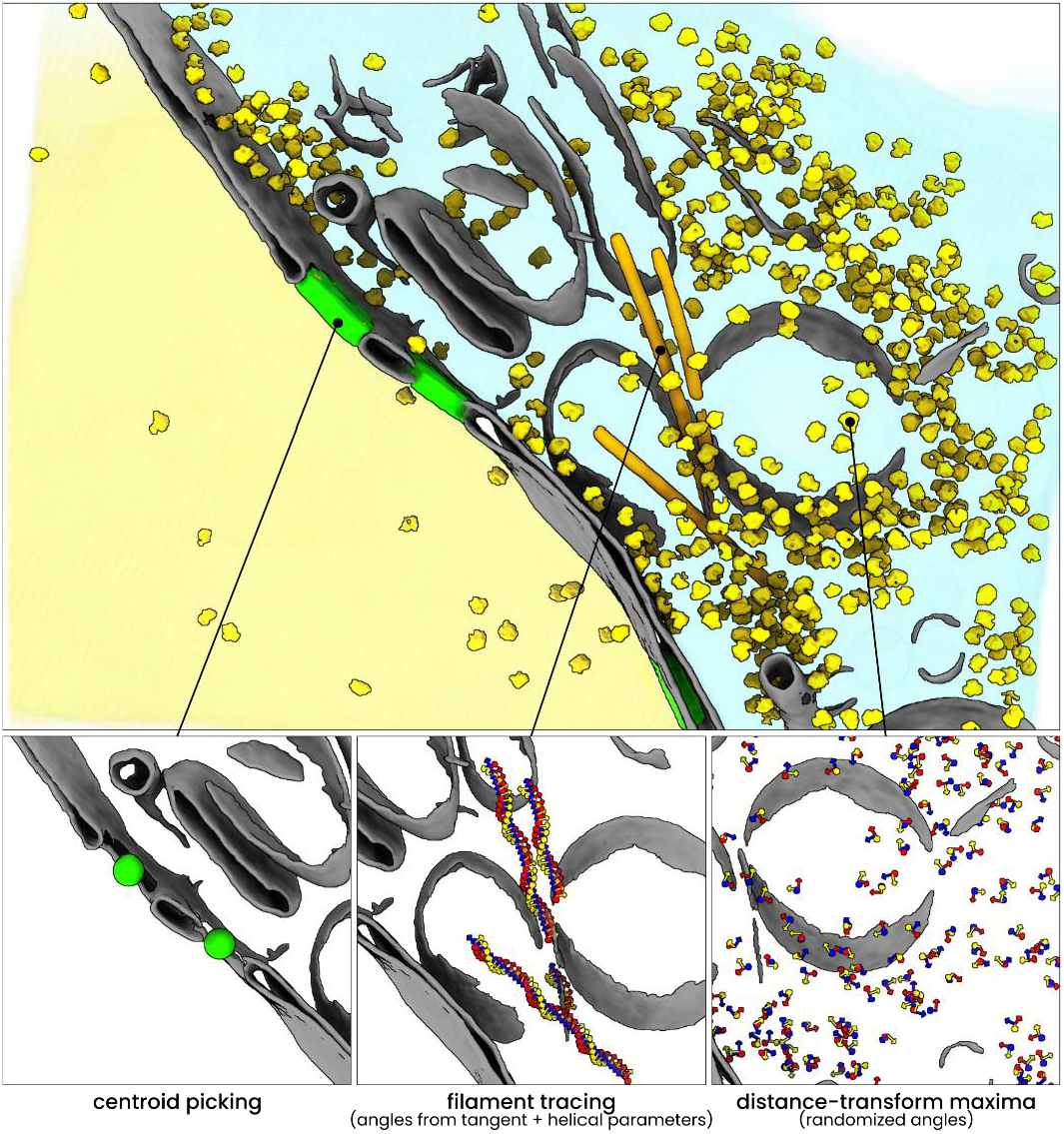
Segmentation-based particle picking. Particle coordinates can be generated from semantic segmentations in three different ways (in easymode), demonstrated here for nuclear pore complexes, microtubules, and ribosomes. **Top:** simplified 3D visualization of the *D. discoideum* tomogram shown in main text Figure 1C-D, showing cytoplasm (cyan), nucleus (yellow), membranes (grey), nuclear pore complexes (green), microtubule (orange), and ribosome (yellow) easymode segmentation outputs. **Bottom:** particle sets visualized using ArtiaX. From left to right the panels show examples the different picking modes. Common to all modes is that segmentation volumes are first thresholded and dust-filtered (i.e., connected areas with a volume below some user-specified threshold are discarded). The modes are: **i) centroid picking:** nuclear pore complex coordinates generated by calculating the centroid coordinates of the green isosurfaces; **ii) filament tracing:** microtubule coordinates generated by fitting a spline to the segmented filaments, then placing coordinates using user-defined helical parameters (optional). The tilt and in-plane angle (rlnAngleTilt, rlnAnglePsi) are derived from the tangent to the spline; **iii) distance-transform maxima:** ribosome coordinates generated by calculating the distance transform of the thresholded segmentation (converting binary voxel values into a value corresponding to the distance to the nearest zero in the binary volume). Coordinates are then placed at the local maxima of this transform, using a user-defined minimum particle spacing. When segmentations of multiple particles may overlap, the distance-transform method is more suitable; when particles are isolated but non-globular, the centroid picking method is generally more suitable.

**Supplementary Figure 2:**
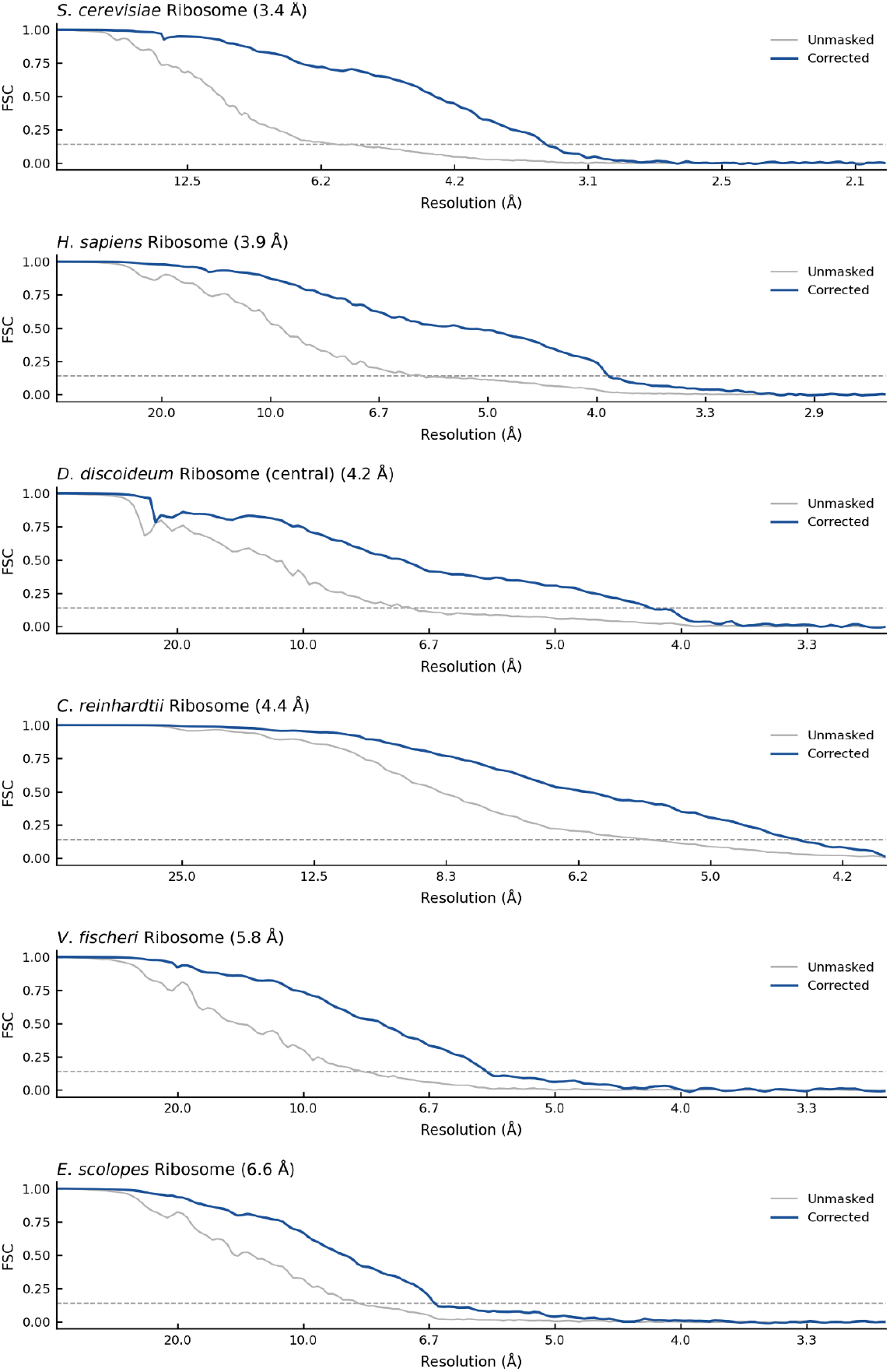
FSC plots for the ribosome maps in Figure 1A. The resolution is reported at the value determined by M after refinement, using a threshold of 0.143 for the corrected half maps. Note that for the *D. discoideum* ribosome, we report the resolution of the improved map based on particles central to the lamella only, as described in main text Figure 3.

**Supplementary Figure 3:**
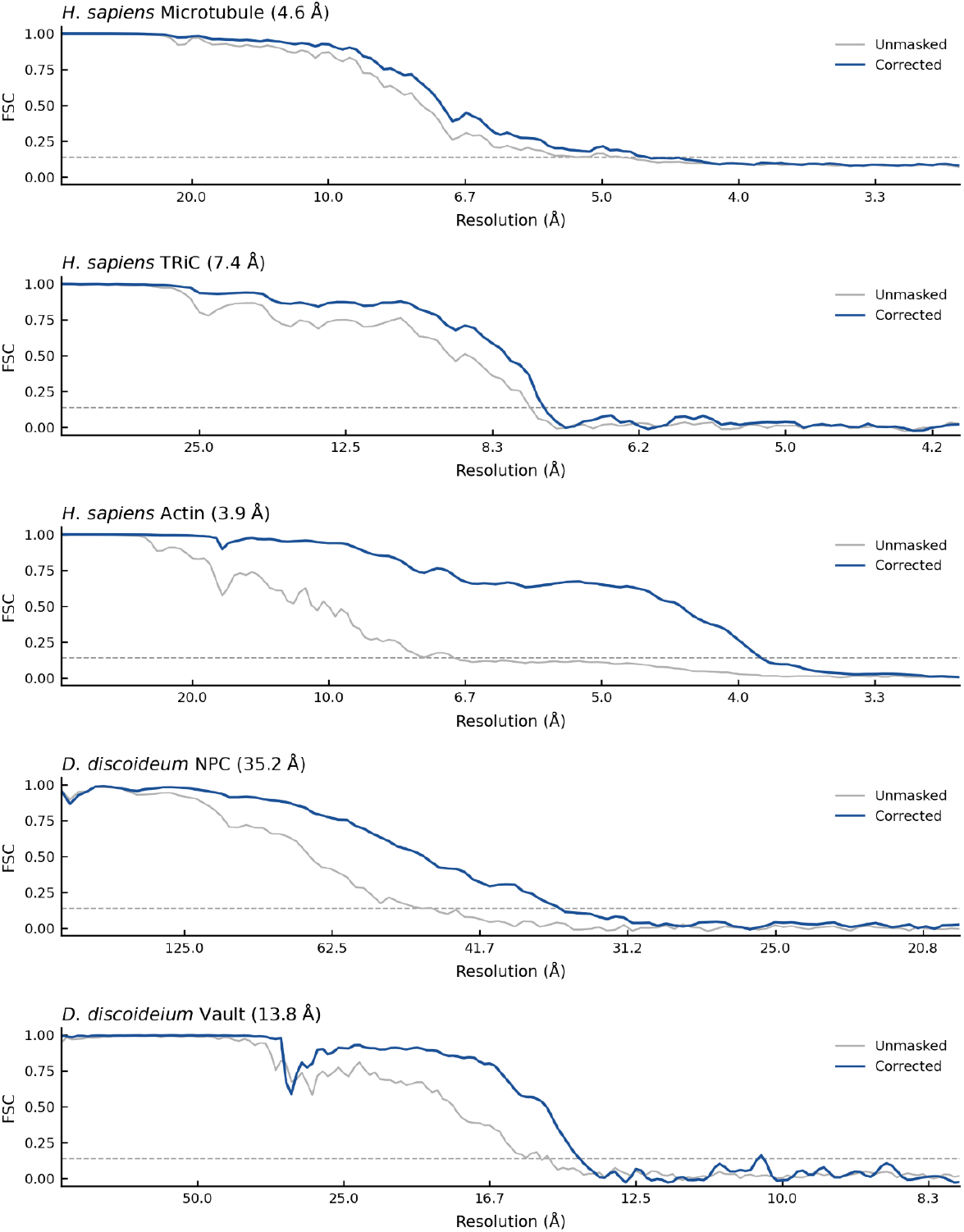
FSC plots for the microtubule, TRiC, actin, NPC, and vault maps in Figure 1. For all maps except the NPC, the resolution is reported at the value determined by M after refinement, using a threshold of 0.143 for the corrected half maps. For the NPC we use the FSC values generated by RELION5 PostProcessing.

**Supplementary Figure 4:**
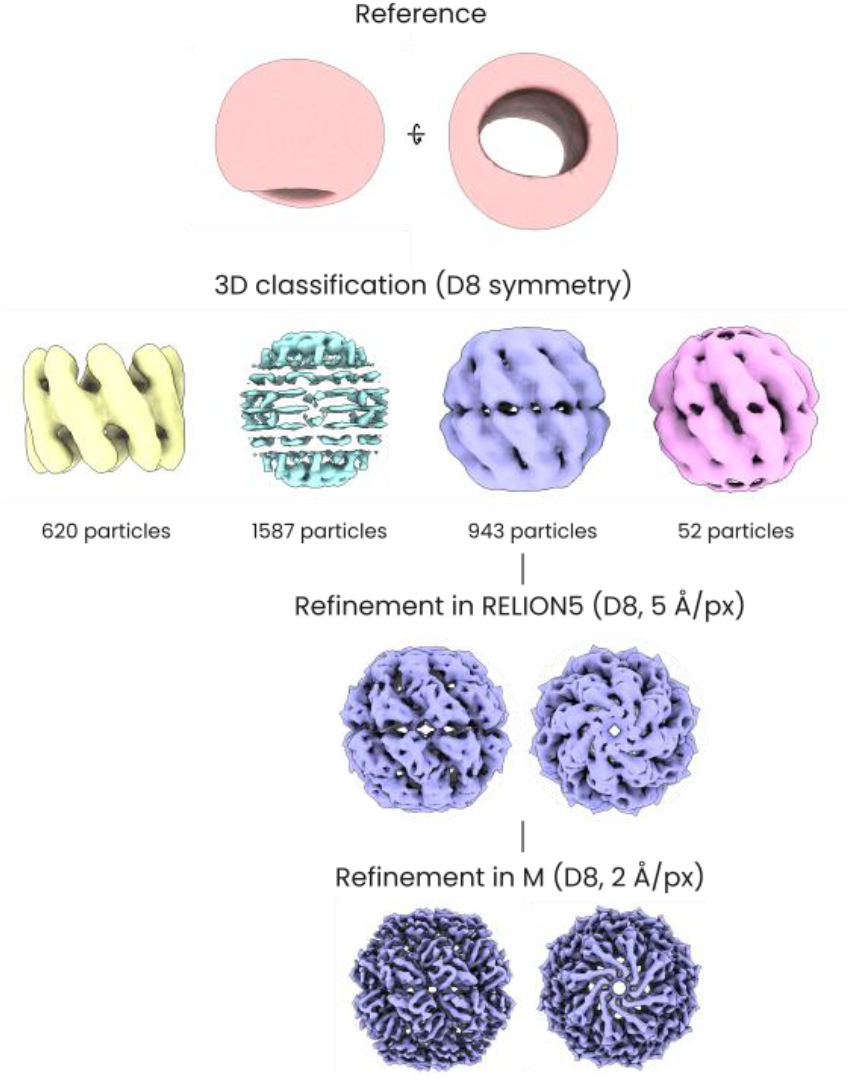
Classification of the TRiC particle set. Subtomogram averaging of TRiC started with a 3D classification (D8 symmetry, 4 classes) in Relion5, using a 80 A low pass filtered map (EMD-18927^79^) as the reference. This yielded a map of what appeared to be TRiC in the open configuration (see EMD-18922^79^; 620 particles), a map of mostly high frequency artefactual density (1587 particles), and two maps that resembled TRiC in the closed state (943 and 52 particles). Using the set of 943 closed-state particles we then continued 3D refinement in Relion5 at 5 Å /px and obtained a map at around 10 Å resolution. Further refinement in M yielded the final map at 7.4 Å resolution.

**Supplementary Figure 5:**
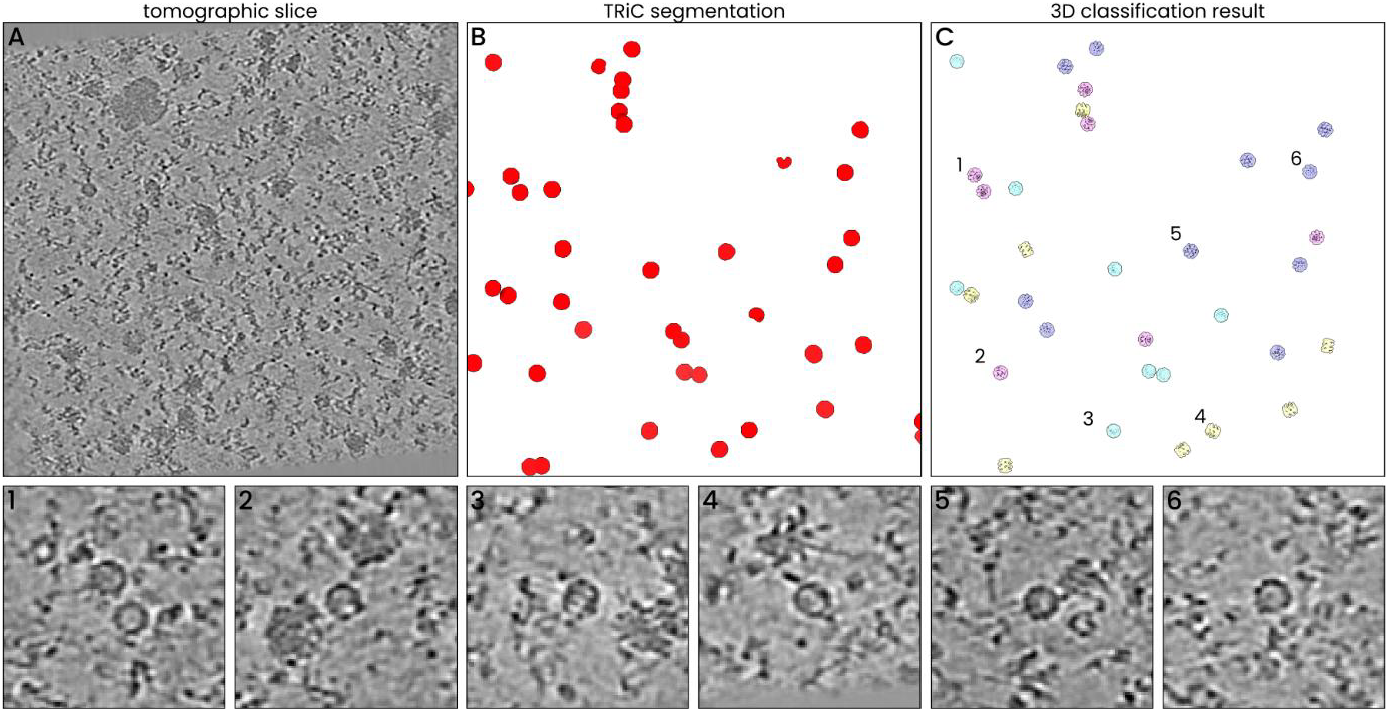
Inspecting 3D classification results of TRiC particles in the original tomogram. **A)** Tomographic slice of an exemplary tomogram. **B)** 3D visualization of the TRiC segmentation output for this tomogram. **C)** Subtomogram averages from 3D classification mapped back into the tomogram volume. The classes are colour-coded in the same way as in Supplementary Figure 4; particles in yellow, cyan, and pink classes were discarded, lavender particles (e.g. those marked 5 and 6) were used for the final map. Examples of these particles in the original tomogram are displayed below, marked 1 – 6. A significant number of particles that were discarded (1 – 4) nevertheless do appear very similar to particles that were included in the final average (5 – 6) and resemble genuine TRiC particles. On the basis of 3D classification alone, we were thus not able to confidently differentiate between false- and true-positive picks.

**Supplementary Figure 6:**
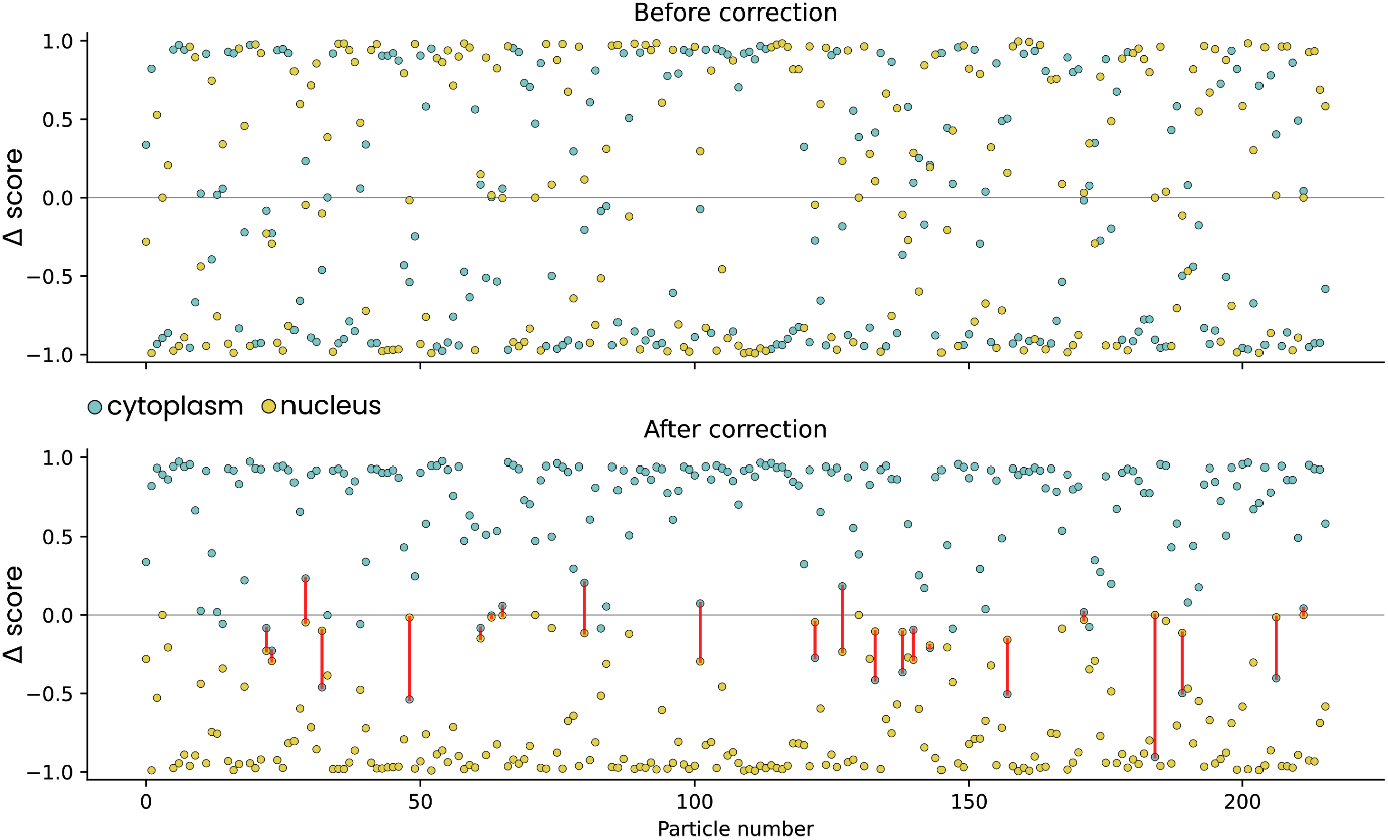
Correcting NPC orientation using segmentations of the cytoplasm and nucleus. After initial 3D alignment, nuclear pore complexes may end up misaligned by 180°, with the distinct cytoplasmic and nuclear sides of the pore facing the wrong way. By segmenting the nucleus and cytoplasm with easymode, it is possible to measure whether a particle is in this misaligned pose. Using the initial pose estimates, we measured the average nucleus and cytoplasm segmentation values within a sphere of 50 nm radius at positions that were offset +25 nm and −25 nm to the particle, relative to its local Z axis (i.e. along the symmetry axis of the pore). These samples take values in a range from 0.0 to 1.0, where 0.0 means that the relevant easymode network output was all zero in that location, and 1.0 means that the network output was maximal in that location. We then calculated a cytoplasm and nucleus ‘Δ score’ for each particle, defined as *c*_*front*_− *c*_*back*_ for the cytoplasm and *n*_*front*_ − *n*_*back*_ for the nucleus (with *c* and *n* indicating the values sampled from the cytoplasm and nucleus segmentations, respectively). For an NPC in the correct pose, these scores should be approximately +1.0 and −1.0; in the incorrect pose, they would be swapped: −1.0 for the cytoplasm and +1.0 for the nucleus, in principle. Based on these scores, we found that the initial pose estimate was flipped for about half of the particles (**top**). The second plot (**bottom**) shows the score distribution after flipping particles based on just the sign of the nucleus score. Some particles were ambiguous – we discarded particles with both a cytoplasm score below 0.3 and a nucleus score above −0.3 (marked in red). If scores are far from the ideal values of +1.0 and −1.0, this can be due to multiple reasons: individual NPCs may be positioned close to the boundaries of the lamella, and instead of surrounded by cytoplasm and nucleus would be surrounded by ‘void’ instead; cytoplasm segmentation averages may also be lower due to the presence of other compartments in front of the NPC (a mitochondrion or vesicle, for example); a third reason may just be that the segmentation output is not perfect.

**Supplementary Figure 7:**
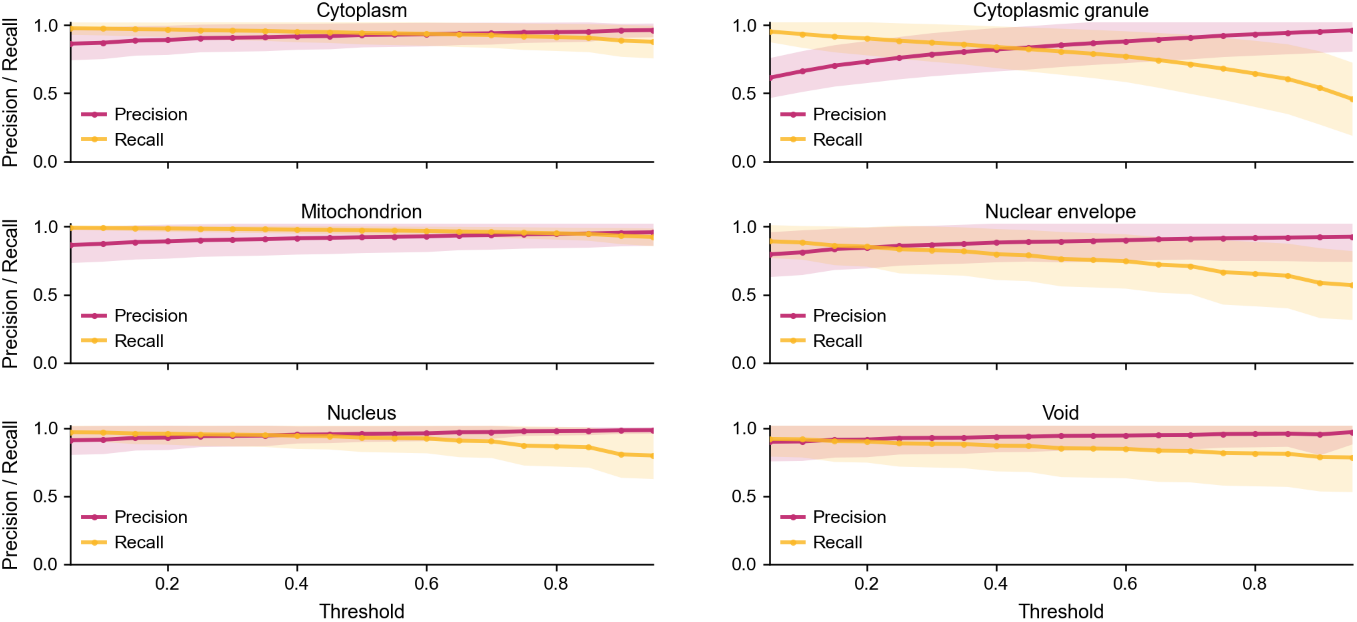
Precision, recall, and DICE scores as a function of threshold value. Precision, recall, and DICE scores as a function of threshold value. The values discussed in the main text were measured using a threshold of 0.5 (with the output range being 0.0 – 1.0, but can be calculated for any threshold value. Lines show the mean and shaded bands ±1 standard deviation across test samples (n = 87 cytoplasm, 42 cytoplasmic granule, 103 mitochondrion, 43 nuclear envelope, 47 nucleus, 63 void). Depending on the application, users may want to optimize for either precision or recall. Generally, lowering the threshold increases recall at the cost of decreased precision.

**Supplementary Figure 8:**
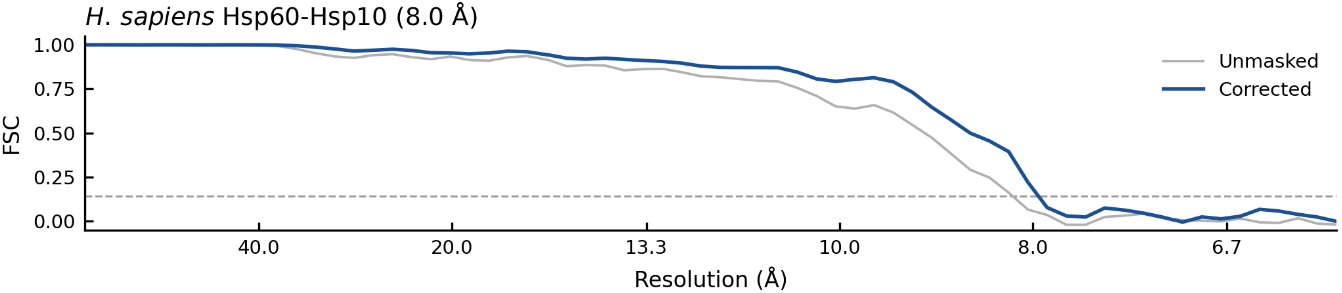
FSC plot for the Hsp60-Hsp10 map in Figure 3D. The resolution is reported at the value determined by M after refinement, using a threshold of 0.143 for the corrected half maps.

**Supplementary Figure 9:**
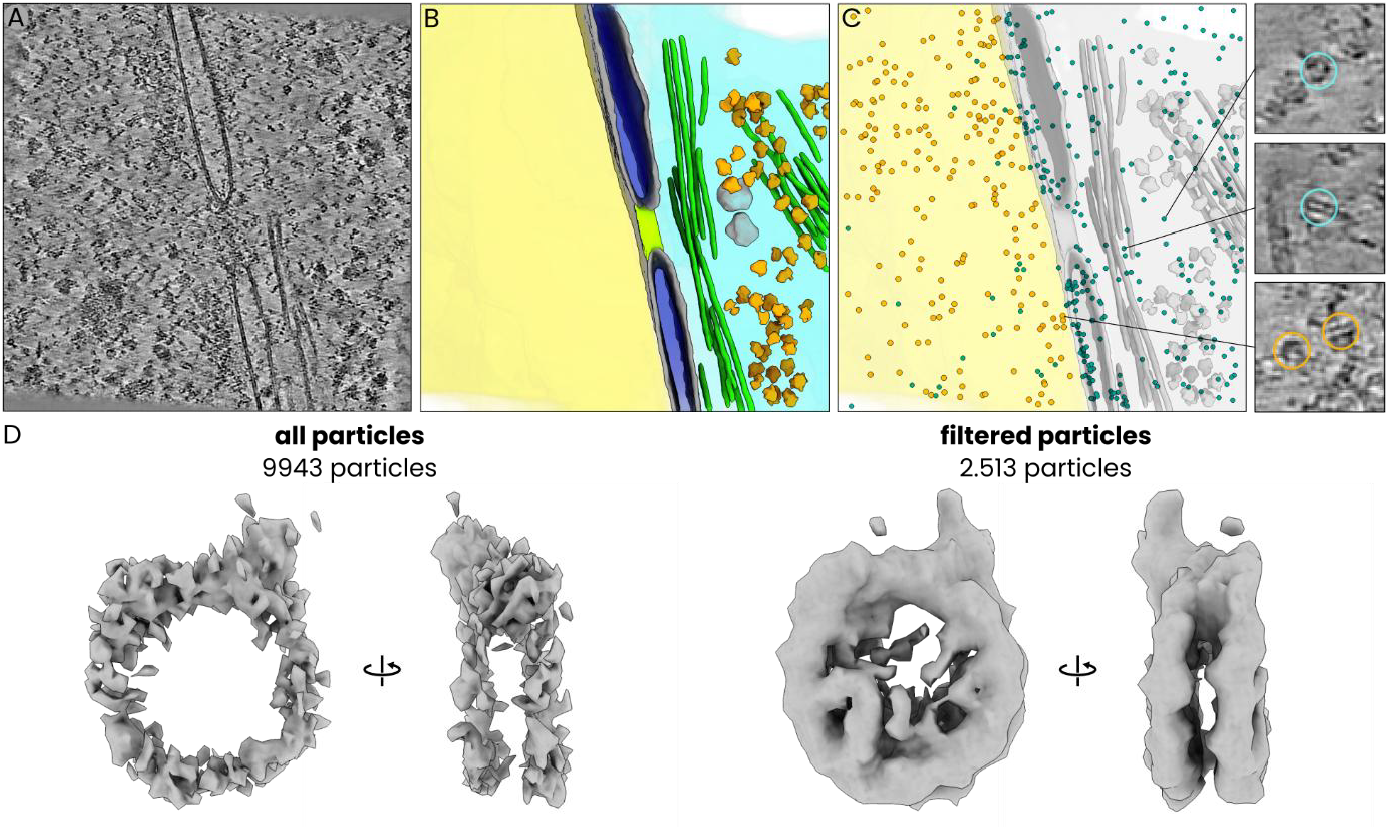
Filtering template matching-derived candidate nucleosome particles using the easymode general nucleus segmentation network. **A)** Tomographic slice of a human (HeLa) cell (same dataset as discussed around main Figure 4), **B)** Visualization of nucleus (yellow), cytoplasm (cyan), membranes (grey), nuclear pore complex (lime), intermediate filaments (green), cytoplasmic granules (light grey), and ribosomes (orange), segmented with easymode. **C)** The location of candidate nucleosomes determined via template matching in PyTom^33^, with particles detected outside of the nucleus indicated in cyan and putative true nucleosomes, detected in the nucleus, in orange. The inset shows examples of two cytoplasmic candidate particles (top, middle) and two presumably true positive nucleosome particles found in the nucleus (bottom). **D)** Subtomogram averages of the filtered and full particle sets, result after 3D refinement and one round of 3D classification. The maps are displayed at the same isosurface level.

**Supplementary Figure 10:**
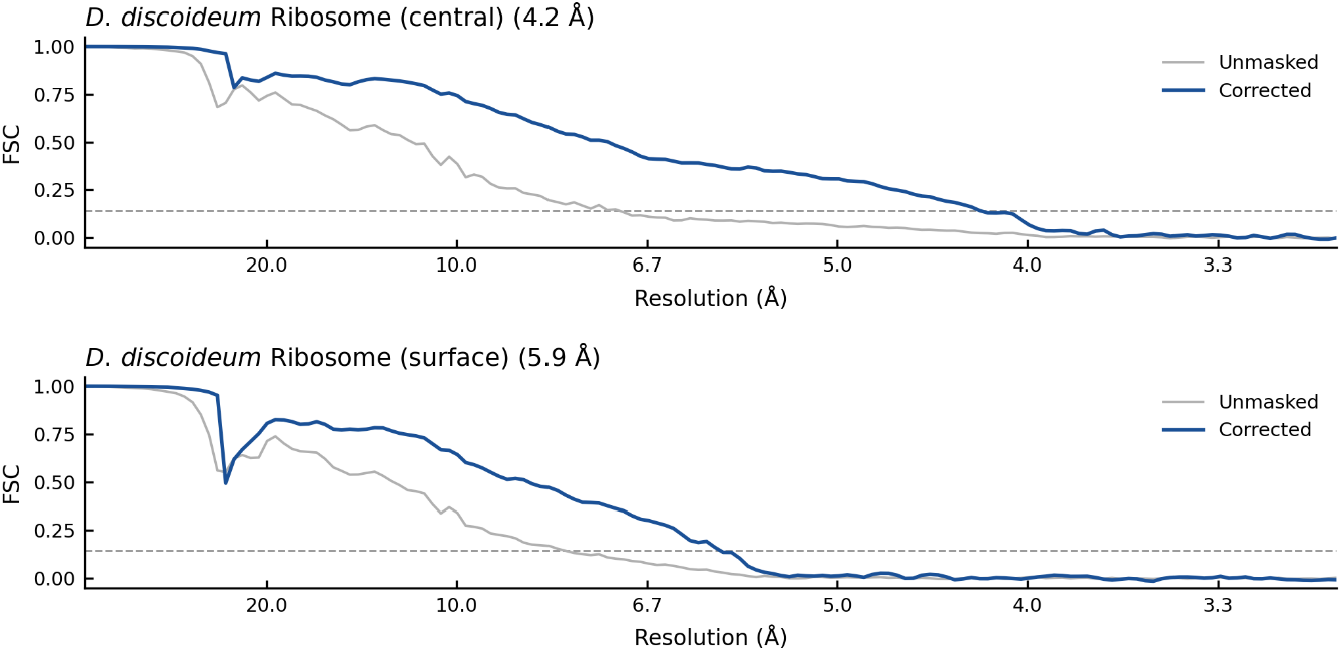
FSC plot for the two *D. discoideum* ribosome maps in Figure 3H. The resolution is reported at the value determined by M after refinement, using a threshold of 0.143 for the corrected half maps.

**Supplementary Figure 11:**
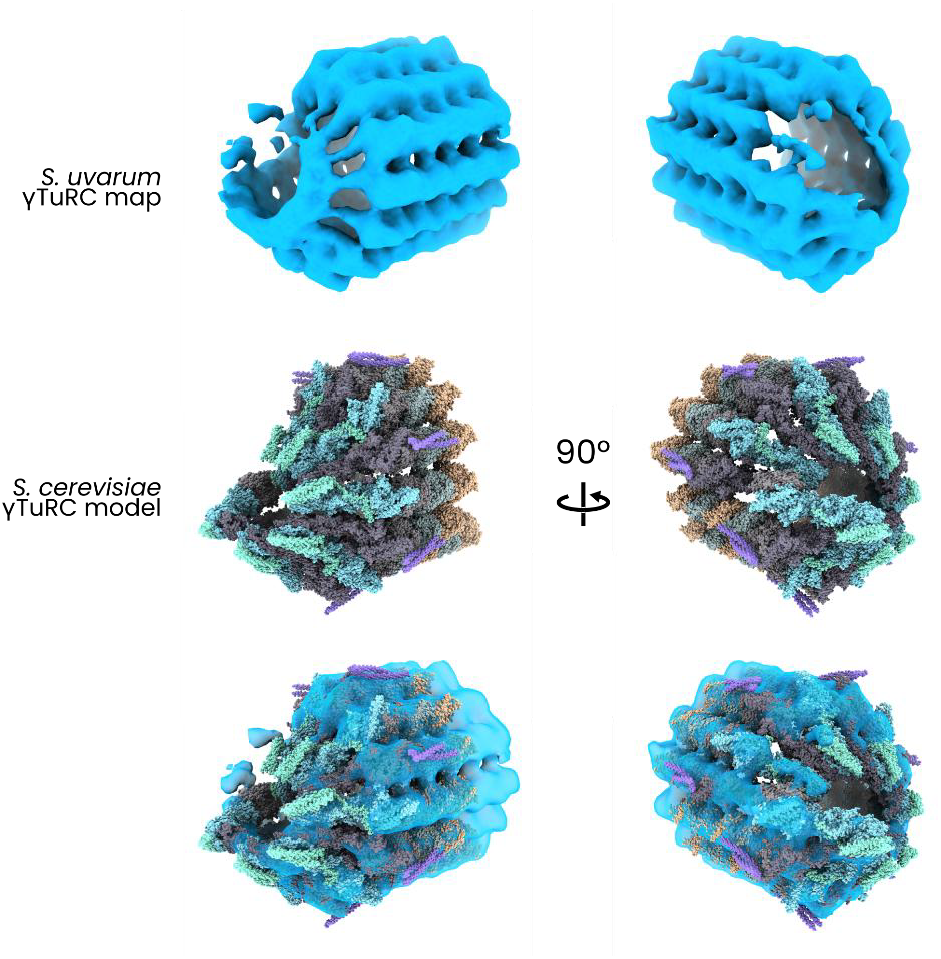
*S. cerevisiae* γTuRC model fit into the *S. uvarum* subtomogram average. Top: two views of the *S. uvarum* γTuRC subtomogram average at 23.6 Å resolution. Middle: corresponding views of a model of the *S. cerevisiae* γTuRC (PDB 8QV2^82^). Below: S. cerevisiae γTuRC model fitted into the *S. uvarum* subtomogram average.

**Supplementary Figure 12:**
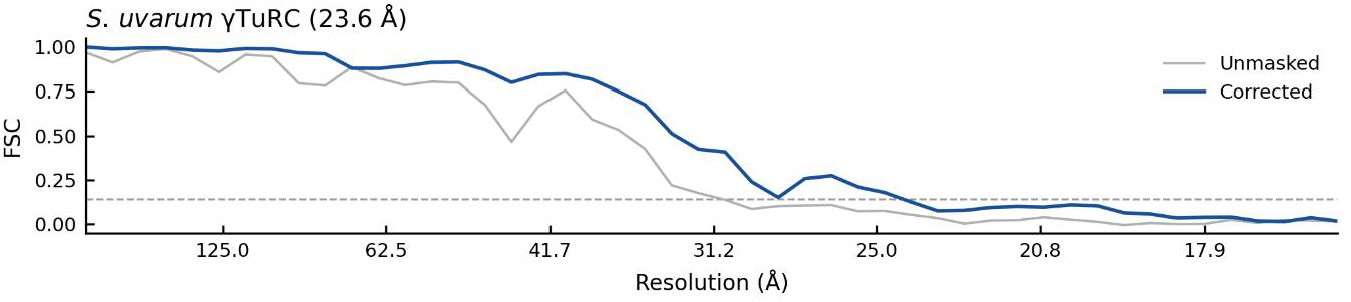
FSC plot for the γTuRC map in Figure 3K. The resolution is reported at the value determined by M after refinement, using a threshold of 0.143 for the corrected half maps.

**Supplementary Figure 13:**
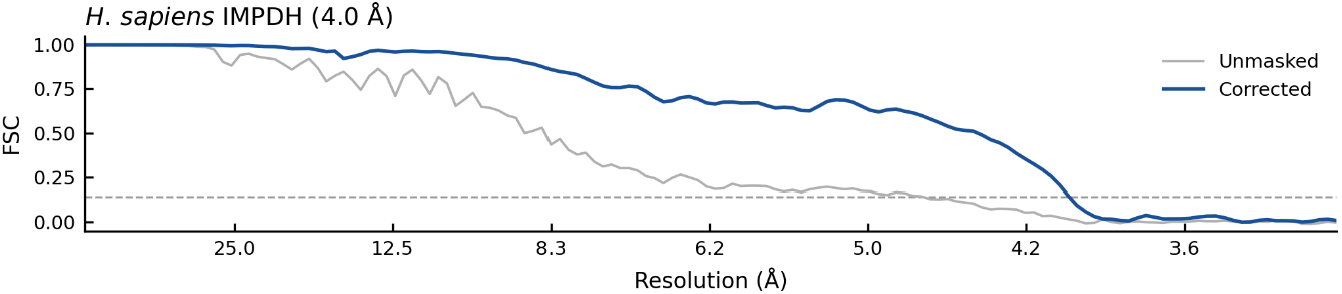
FSC plot for the IMPDH map in Figure 4. The resolution is reported at the value determined by M after refinement, using a threshold of 0.143 for the corrected half maps.

**Supplementary Figure 14:**
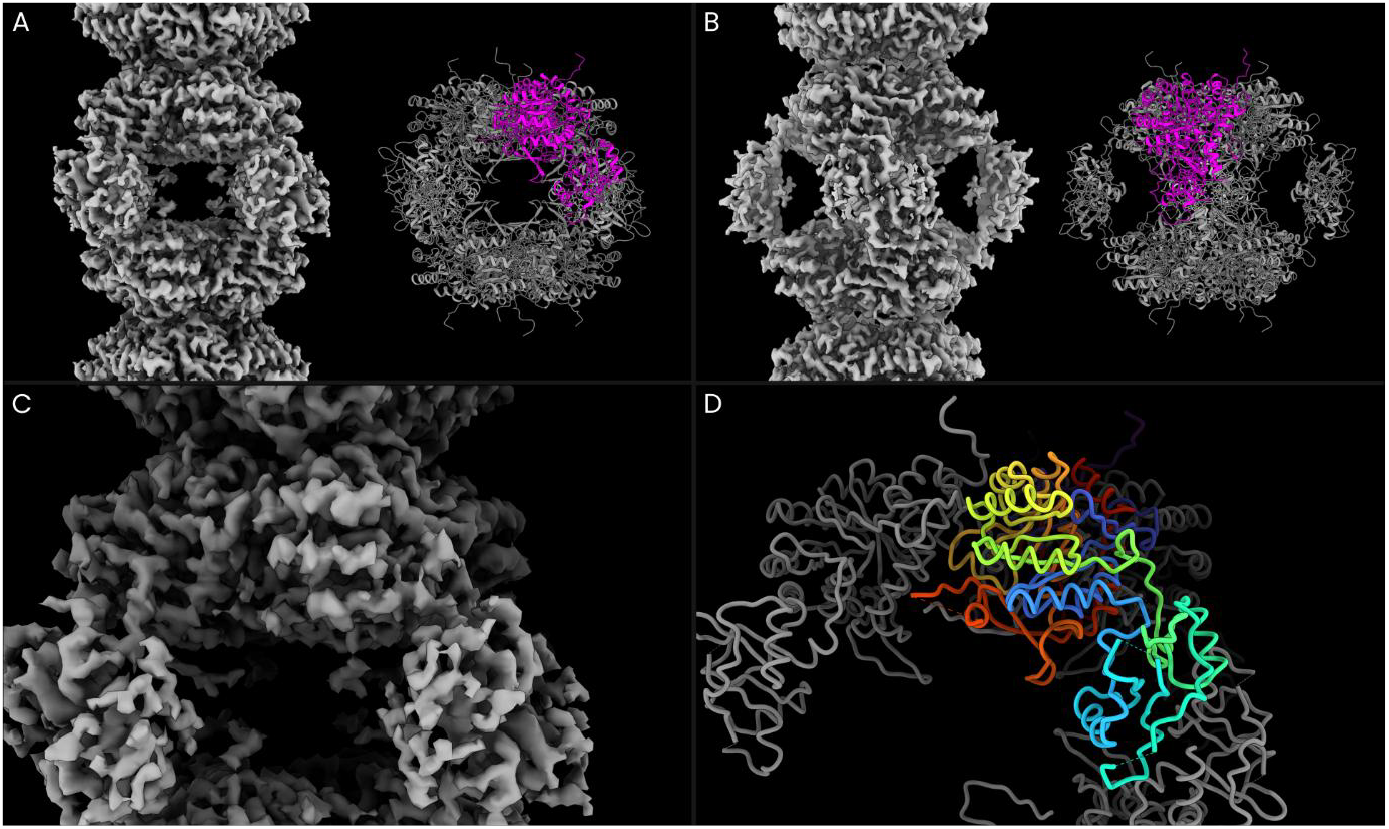
Comparison of the IMPDH subtomogram average to a model of octameric IMPDH1. **A)** Subtomogram average of IMPDH at 4.0 Å resolution (left) alongside a model of an IMPDH isoform 1 octamer in the open state (PDB 7RES^84^), with one monomer highlighted in magenta. **B)** The same map and model as in A, rotated 45° along the filament axis. **C)** A magnified view of the map, centred on the catalytic domain of one monomer. **D)** The same view as in C, showing a single monomer (rainbow) and its immediate neighbours within the same octamer.

**Supplementary Figure 15:**
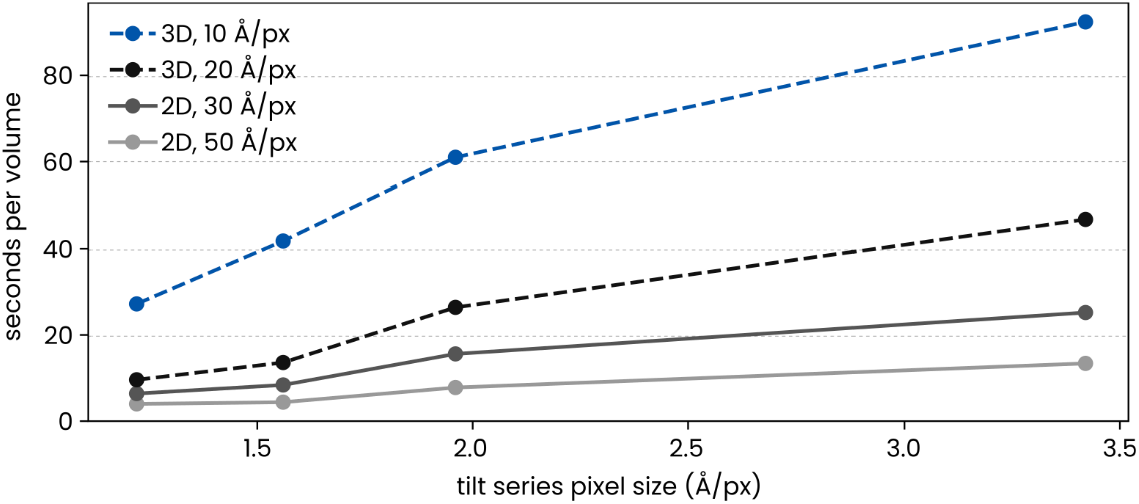
Measurement of the processing speed of easymode networks. Easymode uses a number of different network architectures and pixel sizes at inference for different features. Processing speed therefore differs between different features, and is also strongly dependent in the size of the input volume. The plot shows the processing cost of segmentation in seconds per tomogram volume per GPU for four different networks/pixel sizes and for four example datasets. These datasets span the range from relatively high (1.22 Å /px) to low magnification (3.42 Å /px). The 2D network at 50 Å /px (used for, e.g., cytoplasm, nucleus, mitochondrion) is generally fastest, followed by the 2D network at 30 Å /px (used for, e.g., the nuclear envelope, void). The 3D network is slower. At 20 Å /px (used for cytoplasmic granules, mitochondrial granules), segmentation of a single tomogram costs between 10 – 50 seconds. At 10 Å /px, which is the scale used for most features (including ribosomes, microtubules, vault complexes, actin, intermediate filaments, TRiC, membrane), processing cost for a single volume is ∼25 seconds for the high magnification dataset and ∼90 seconds for the low magnification dataset. When using multiple GPUs individual volumes are processed in parallel so that overall processing time reduces linearly with the number of GPUs available.

**Supplementary Figure 16:**
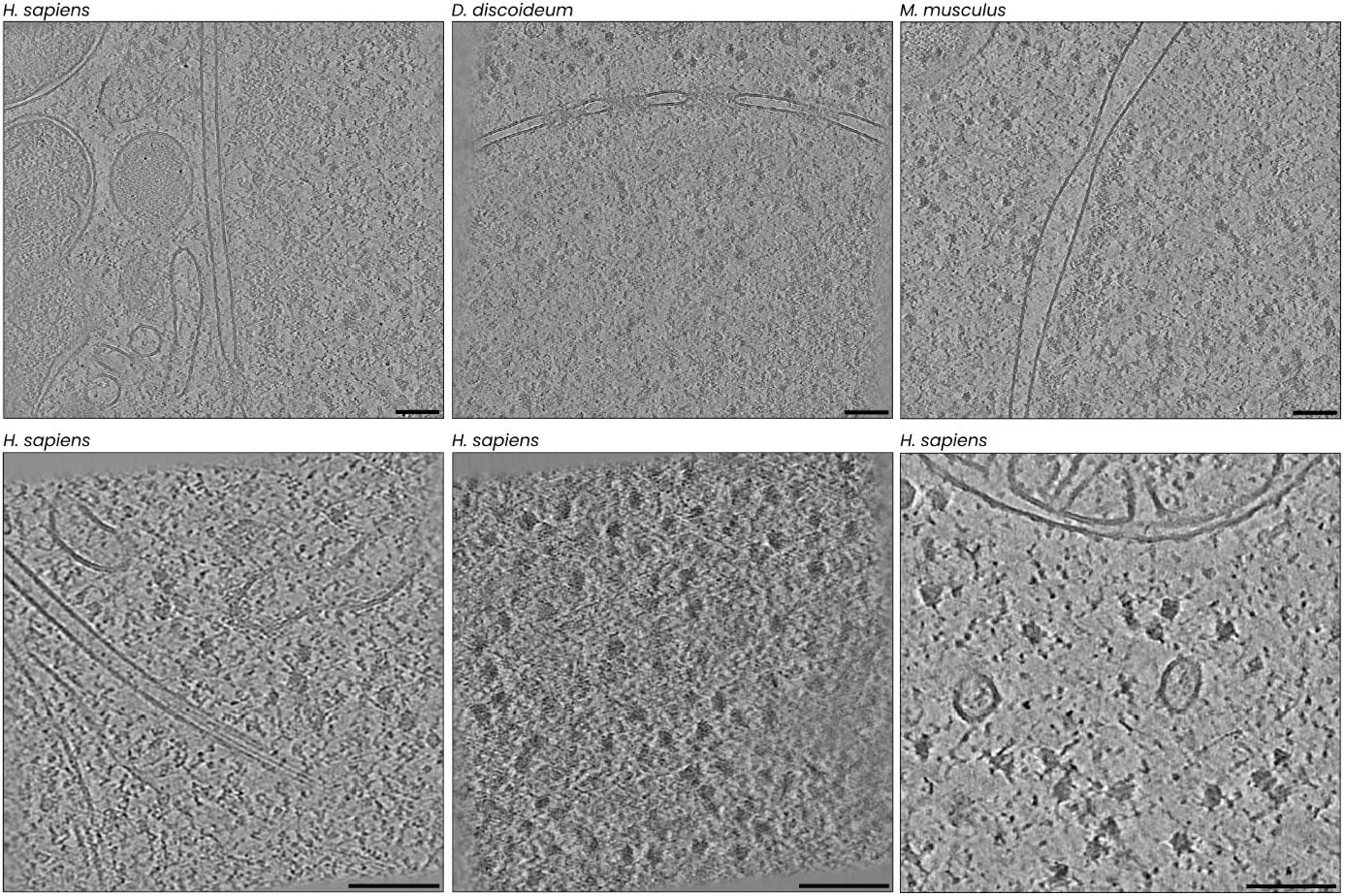
Sources of variation in cryo-ET datasets. Top: tomographic slices of FIB-milled lamellae of different species; *H. sapiens* (macrophage, EMPIAR-12457), *D. discoideum* (EMPIAR-11845), and *M. musculus* (embryonic stem cell, EMPIAR-12460) cells, acquired with similar acquisition parameters (pixel sizes 2.2 – 2.7 Å /px, defocus 2.8 – 3.2 μm). Bottom: tomographic slices of the same species, acquired with more varied acquisition parameters: 1.19 Å /px with 4.7 μm defocus (EMPIAR-11538; left), 1.19 Å /px with 1.4 μm defocus (EMPIAR-11538; middle), and 3.43 Å /px with 3.1 μm defocus (EMPIAR-11561; right). Scale bars are 100 nm. Based on our experience working with the easymode training collection, we find that, overall, the acquisition parameters (including pixel size, defocus, dose) are a larger contributor to variation between tomograms than differences between different (eukaryotic) species or cell types.

